# Deciphering epileptogenic and activity-dependent gene programs in the human brain

**DOI:** 10.64898/2026.02.18.706645

**Authors:** Qianyu Lin, Chris Kang, Andre M. Xavier, Irene Sanchez-Martin, Ashesh D. Mehta, Lucas Cheadle

**Affiliations:** Cold Spring Harbor Laboratory, Cold Spring Harbor, NY, 11740, USA; Feinstein Institutes for Medical Research, Northwell Health, Manhasset, NY 11030, USA; Howard Hughes Medical Institute, Cold Spring Harbor, NY 11740, USA

## Abstract

Neuronal activity is fundamental to brain function, yet chronically elevated activity underlies neurological disorders such as drug-resistant epilepsy (DRE). In animal models, activity induces defined transcriptional programs within activated neurons; however, the nature, cellular specificity, and pathological relevance of such programs in the human brain remain poorly understood. Here, we apply single-nucleus and spatial transcriptomics to epileptogenic, non-epileptogenic, and intraoperatively stimulated non-epileptogenic cortical tissue obtained from individuals with DRE. Across 26 cell types profiled, glutamatergic neurons projecting from cortical layers 2/3, 5, and 6 to intratelencephalic targets exhibit pronounced sensitivity to the epileptogenic microenvironment, inducing shared immediate-early genes alongside cell-type-specific programs linked to synaptic remodeling and cellular stress. Approximately one-third of transcripts enriched in the epileptogenic microenvironment were also induced by acute stimulation, suggesting that a fraction of epilepsy-associated gene expression reflects conserved responses to heightened activity rather than disease-specific programs. While transcripts induced by both epileptogenic and acute activity converged upon immediate-early genes and heat-shock proteins, only acute stimulation triggered a rapid, multicellular induction of transcripts involved in mitochondrial ATP synthesis. This divergence suggests that neurons within epileptogenic cortex may be unable to mount appropriate metabolic adaptations to sustained energetic demands. In parallel, both microglia and circulating CD14+ monocytes exhibit signs of immune activation in epilepsy, suggesting myeloid-driven inflammatory rewiring that extends beyond the brain. Together, these findings illuminate human activity-dependent gene programs and reveal signatures of neuronal vulnerability and inflammation in DRE.

## Introduction

Neuronal activity is the language of the brain, orchestrating electrochemical communication between neurons and contributing to all brain functions including sensory processing, cognition, and behavior. For example, critical roles for neuronal activity include shaping the assembly and refinement of neural circuits during brain development^1–8^ and facilitating learning and memory in mature organisms^9–12^. At the mechanistic level, work in animal models has shown that activity governs brain function in part by inducing defined transcriptional programs in the nuclei of synaptically activated neurons^13–15^. These programs include an initial wave of immediate-<u>e</u>arly <u>g</u>enes (IEGs) such as the transcription factors *Fos*, *Npas4*, and *Egr1*^16–19^ which, upon their induction, bind defined motifs across the genome to drive the transcription of a second wave of late-<u>r</u>esponse <u>g</u>enes (LRGs). These LRGs encode effector proteins with defined roles in neuronal and synaptic function such as the neurotrophic factor BDNF^20^, the secreted neuropeptide Scg2^21^, and the cytokine receptor Fn14^22–24^. As core transcriptional regulators, activity-dependent IEGs tend to be shared across brain cell types, while LRGs are largely non-overlapping, reflecting the role of activity in shaping neuronal function in a precise cell-type-specific manner. While mechanistic insights into activity-dependent gene expression have been predominantly derived from studies in animals, a handful of studies in iPSC-derived human neuronal cultures suggest partial evolutionary conservation of these activity-dependent gene programs^25–30^, consistent with epigenomic analyses of healthy human brain tissue^31,32^. However, the nature, cellular specificity, and pathological relevance of activity-dependent gene expression in the intact human brain remain to be defined.

While activity is central to healthy brain function, aberrantly heightened activity underlies epilepsy, a common and debilitating disorder characterized by seizures arising from imbalances in excitation and inhibition^33^. In focal epilepsy, seizures are generated by one or more epileptogenic areas within the cortex. Unlike relatively healthy cortical tissue, this <u>e</u>pileptogenic <u>m</u>icro<u>e</u>nvironment (eME) is characterized by commonly recurring seizures, or electrical spikes reflecting glutamatergic neuronal firing, interspersed with interictal discharges^34,35^. Over time, the eME accumulates not only signatures of heightened neuronal activity but also signs of localized neuroinflammation, blood-brain-barrier compromise, and related pathological signatures^36^. While epilepsy symptoms can be effectively managed in some cases, around 1/3 of patients do not respond to currently available antiepileptic treatments^35^. For these patients with drug-resistant epilepsy (DRE), surgical intervention is frequently required but these procedures are highly invasive and, in many cases, insufficient to fully resolve symptoms^37^. Thus, developing new, less invasive approaches for treating DRE is a key unmet goal of biomedical research.

Insights into the cellular and molecular mechanisms underlying the pathophysiology of DRE could identify novel targets for therapeutic intervention. While several studies have employed single-cell RNA sequencing (scRNAseq) approaches to identify molecular disease signatures in the epileptic brain, these experiments were subject to three important caveats^38–42^. (1) These studies compare cortical tissue collected from individuals with DRE to healthy tissue collected from a separate cohort. This approach introduces additional experimental variables beyond disease state such as age, sex, and anatomical region analyzed. Even when these factors are aligned, inter-individual variability could hamper interpretation. (2) Tissue samples collected from individuals with DRE commonly include both epileptogenic and non-epileptogenic tissue, again complicating the interpretation of comparative transcriptomic experiments. (3) As the epileptogenic focus is defined by seizure activity, transcripts enriched within epileptogenic cortex compared to healthy cortex likely include both disease-associated transcripts and transcripts induced as part of a cell’s normal response to heightened activity. Disambiguating which transcripts enriched in epileptogenic cortex represent disease signatures and which represent normal cellular responses to activity could shed light on the factors most likely to promote disease, while also unveiling activity-dependent gene programs in non-epileptogenic human brain tissue.

To shed light on the molecular underpinnings of DRE while addressing these caveats, we performed single-nucleus RNA sequencing (snRNAseq) on anatomically matched epileptogenic and non-epileptogenic cortical brain tissue collected from the same individuals undergoing surgical resection for DRE. To disambiguate disease- and activity-regulated transcripts, a portion of non-epileptogenic cortex was electrically stimulated thirty minutes prior to resection in a subset of patients, then collected and analyzed alongside the other samples. By profiling epileptogenic, non-epileptogenic, and acutely stimulated non-epileptogenic tissue from the same individuals, our transcriptomic analysis spanning 26 distinct cell types uncovers the impact of acute stimulation and epileptogenic activity on neuronal and non-neuronal gene expression. These data represent an unbiased transcriptomic atlas of disease-related and activity-dependent gene programs in the human brain, revealing key insights into cellular vulnerability and resilience in DRE.

## Results

### Single-cell transcriptomics reveals cell-type-specific responses to epileptogenic activity and acute stimulation in the human brain

To characterize gene signatures associated with the <u>eME</u>, we collected brain tissue from the cortices of six individuals undergoing surgical resection for DRE. Tissue was collected from two cortical regions defined by presurgical and intraoperative mapping: (1) the seizure-onset zone (i.e. the epileptogenic focal region) and (2) an anatomically matched non-epileptogenic zone (i.e. a non-focal region devoid of seizure activity)(**Extended Data Fig. 1a-f**). To aid in the disambiguation of disease- and normal neuronal activity-dependent gene signatures, we also collected non-focal tissue subjected to acute stimulation *in vivo* thirty minutes prior to resection. In brief, a handheld bipolar ball tip probe was used to apply focal biphasic stimulation with a maximum current of 10 mA, corresponding to a 20 mA differential. Long-train bipolar stimulation was delivered at a frequency of 60 Hz with a pulse width of 500 microseconds (μs) and a train duration of 5 seconds. All tissue types were immediately flash frozen and single nuclei were isolated and subjected to RNA sequencing using the 10X platform (**Fig. 1a**). In total, 19 samples were sequenced across six patients, including eight focal, seven non-focal, and four acutely stimulated non-focal regions (**Fig. 1b**). Following next generation sequencing, we mapped reads to the human genome then applied a stringent quality control pipeline including removal of ambient RNA, putative doublets, and low-quality or dead cells to obtain a final dataset including 82,604 high-quality nuclei with at least 1000 UMIs per cell. Guided by the Allen Brain Atlas^43,44^, we annotated 26 distinct cell types spanning glutamatergic and GABAergic neurons, glia, vascular cells, and immune cells (**Fig. 1c,d and Extended data Fig. 1g-i**). In parallel, we subjected a subset of the focal (i.e. epileptogenic) and non-focal (i.e. non-epileptogenic) samples (6 focal, 5 non-focal) to spatial analysis using the Xenium platform, allowing us to spatially map 149,521 segmented cells annotated based upon our snRNAseq results, validating the cortical architecture and cellular composition of the samples *in situ* (**Fig. 1e-g and Extended data Fig. 2**). By profiling diverse cell types across focal, non-focal, and acutely stimulated tissue, this dataset delineates the transcriptomic adaptations of neuronal and non-neuronal human brain cells to the eME and to acute stimulation *in vivo*.

**Figure 1.**
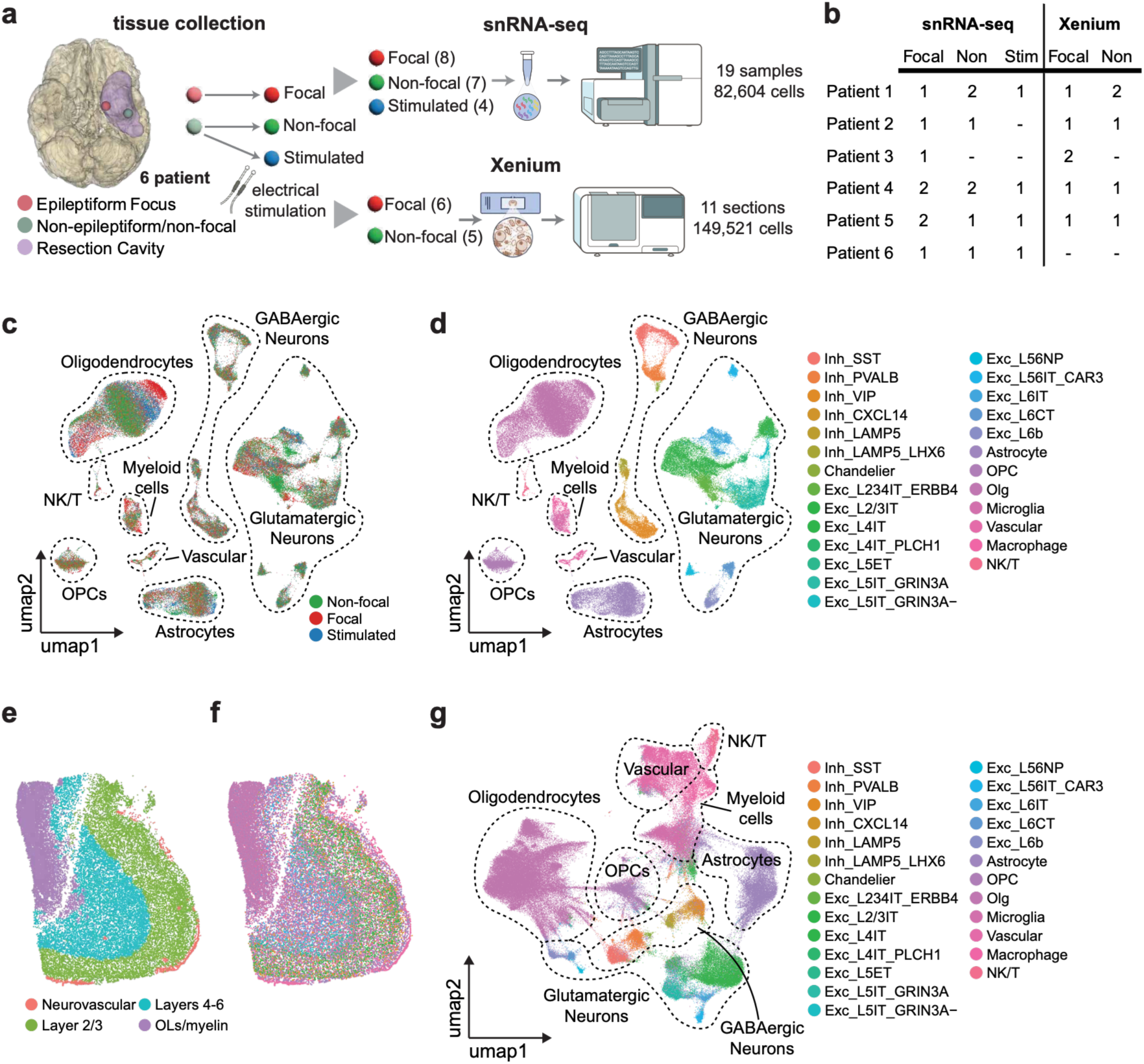
A transcriptomic atlas of epileptogenic and acute activity-dependent gene expression in the human brain. (a) Schematic of the experimental approach to perform multi-modal transcriptomic profiling of freshly resected cortical tissue from individuals with DRE. (b) Table listing the number and types of cortical samples sequenced by patient. (c) Uniform Manifold Approximation and Projection (UMAP) plot of all cells in the dataset colored by sample type, legend in bottom right. (d) UMAP colored by cell type, legend on right. (e) Representative non-epileptogenic cortical section profiled by Xenium, colored by cellular niche. (f) Same as (e) but colored by cell type according to legend in (g). (g) UMAP of Xenium data by cell type.

### Intratelencephalic-projecting (IT) glutamatergic neurons in layers 2/3, 5, and 6 are highly responsive to the eME

To determine how the eME impacts the transcriptomic signatures of cortical cells, we began by performing differential gene expression analysis between focal and non-focal regions using Seurat(v5) with patient identity included as a latent variable to control for inter-individual effects^45^. Genes with a fold-change ≥ log2(1.5) meeting a stringent significance threshold of false discovery rate (FDR) < 0.05 were considered significantly differentially expressed genes (DEGs). This analysis uncovered robust alterations in the transcriptomic profiles of cells within focal compared to non-focal regions, identifying a total of 1,301 and 711 unique transcripts that are upregulated or downregulated, respectively, within the epileptogenic focus. These data reveal substantial transcriptomic differences between anatomically matched regions of epileptogenic and non-epileptogenic cortex from the same brains.

We next asked whether some cell types were more sensitive to the eME than others by comparing the numbers of DEGs identified between focal and non-focal tissue. This analysis uncovered striking heterogeneity in DEG abundance, with some cellular populations exhibiting fewer than 50 DEGs and others exhibiting over 300 (**Fig. 2a**). Among all cell types in the dataset, subpopulations of glutamatergic neurons mounted a stronger transcriptomic response to the eME than GABAergic neurons or glia, with three intratelencephalic (IT) projection neuron subsets accumulating the largest numbers of DEGs: glutamatergic layer 2/3-IT (Exc_L2/3IT; 382 DEGs), GRIN3A+ glutamatergic layer 5 IT (Exc_L5IT_GRIN3A+; 630 DEGs), and glutamatergic layer 6 IT (Exc_L6IT; 318 DEGs) neurons. Conversely, glutamatergic neurons that do not project intratelencephalically, such as extratelencephalically projecting layer 5 (Exc_L5ET; 3 DEGs) and near-projecting layer 5/6 (Exc_L5/6IT_CAR3; 40 DEGs) neurons, exhibited few transcriptomic differences between focal and non-focal regions (**Fig. 2a**). As the number of cells per cluster can influence the detection of DEGs based upon single-cell transcriptomics, we also assessed DEG abundance between cell types after downsampling all populations to 500 cells. Even after downsampling, Exc_L2/3IT, Exc_L5IT_GRIN3A+, and Exc_L6IT neurons exhibited the greatest DEG abundance among all cell types in the dataset (**Extended data Fig. 3a**). These observations suggest that subpopulations of glutamatergic neurons are more responsive at the transcriptomic level to the eME than GABAergic neurons or glia, and that, among glutamatergic neurons, cellular vulnerability is more closely linked to neuronal projection pattern than to laminar position.

**Figure 2.**
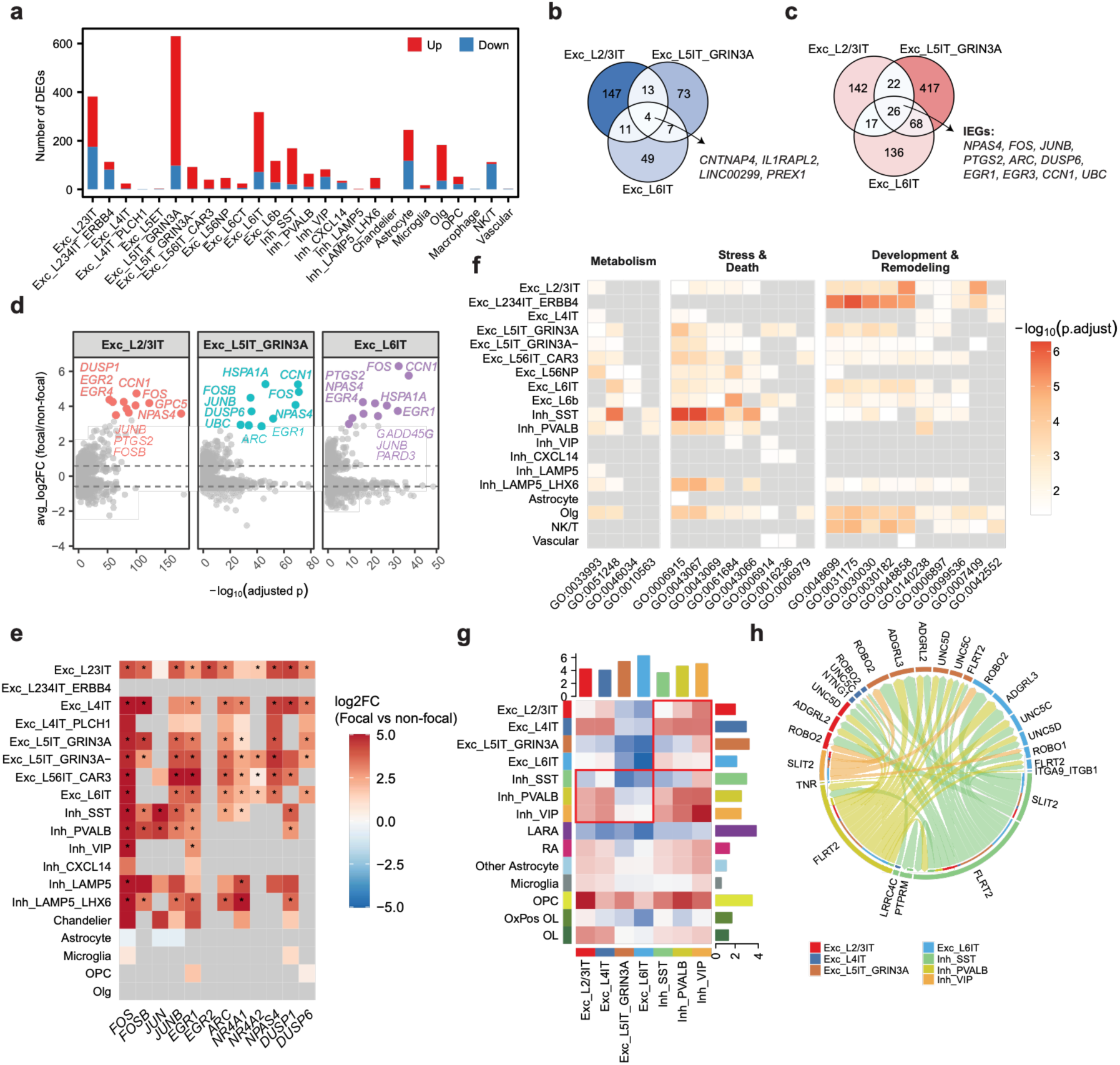
Epileptogenic activity induces shared IEGs and divergent effector programs in intratelencephalic-projecting glutamatergic neurons. (a) Abundance of differentially expressed genes (DEGs) between focal and non-focal regions, thresholded at FDR < 0.05, |log2FC| > log2(1.5). Red, upregulated in focal region; blue, downregulated in focal region. (b) Venn diagram of focally downregulated DEGs shared among Exc_L2/3IT, Exc_L5IT_GRIN3A, and Exc_L6IT neurons. (c) Venn diagram of focally upregulated DEGs shared among Exc_L2/3IT, Exc_L5IT_GRIN3A, and Exc_L6IT. A subset of shared immediate early genes (IEGs) is highlighted. (**d)** Volcano plots of DEGs in Exc_L2/3IT, Exc_L5IT_GRIN3A and Exc_L6IT neurons between focal and non-focal conditions. The top 10 upregulated DEGs in each cell type are highlighted in color, with all other genes shown in grey. Dashed lines indicate FDR = 0.05 and |log2FC| = log2(1.5) thresholds. (e) Heatmap demonstrating canonical IEG expression across major cell types between focal and non-focal regions. Colors indicate log2FC according to the scale on the right, where warmer colors are expressed more highly in the focal region. Asterisks denote significant genes with |log2FC| > log2(1.5) and FDR < 0.05, as determined by MAST (Model-based Analysis of Single-cell Transcriptomics) using a generalized linear hurdle model, with patient identity included as a latent variable to control for inter-individual effects. Non-significant genes are shown in gray; only genes with min.pct ≥ 0.1 were tested. (f) Heatmap of Gene Ontology (GO) enrichment across all cell types based on DEGs between focal and non-focal conditions, broken into functional modules. Significantly enriched GO terms (FDR < 0.05) are colored by −log10(FDR), with non-significant terms shown in grey. (g) Heatmap of differential intercellular communication networks between focal and non-focal regions inferred by *CellChat*. Senders, y axis. Receivers, x axis. Red boxes denote putative signaling from and to GABAergic neurons. (h) Chord diagram showing ligand–receptor signaling from GABAergic neurons to IT-projecting glutamatergic neurons inferred by *CellChat* using DEG-mapped interactions (min. pct >10%, |log2FC| > 0.1).

### Gene programs induced in IT-projecting neurons share a strong IEG signature but otherwise diverge toward synaptic remodeling or cellular stress

We next asked whether the transcriptomic changes undergone by the three most strongly affected IT-projecting populations were shared with one another, or whether they were cell-type-specific. Among transcripts that were downregulated in the focal environment, only four (*CNTNAP4*, *IL1RAPL2*, *LINC00299*, and *PREX1*) were shared between cell types (**Fig. 2b**). Conversely, 26 transcripts were upregulated by epileptogenic activity in all three populations, with canonical IEGs (e.g. *NPAS4*, *FOS*, *JUNB, PTGS2, ARC*, *DUSP6*, *EGR1,* and *EGR3)* being heavily represented within this cohort (**Fig. 2c,d**). Consistent with seizure activity leading to hyperexcitation within the eME, IEGs were broadly enriched across most glutamatergic populations within the focal region regardless of projection pattern, while most GABAergic neurons induced a more restricted repertoire of IEGs that included AP-1 transcription factors such as *FOS* and *JUN* but not *NR4A2* or *NPAS4* (**Fig. 2e**). However, the majority of focally enriched genes within IT-projecting cells were non-overlapping, and gene ontology (GO) analysis suggests that their functions differ between neurons in upper and deep layers. For instance, IT-projecting neurons in L2/3 induce transcripts involved in axonogenesis and synaptic remodeling (e.g. *SHTN1*, *ROBO1*) within the eME, while IT neurons in L5 and L6 induce transcripts involved in the cellular stress response (e.g. *HSPA1A*, *FTH1*) and cell death pathways (e.g. *MOAP1*, *CTSD*; **Fig. 2f**). Thus, upper layer IT-projecting neurons may be better equipped to adapt to seizure activity through circuit rewiring, whereas deep layer neurons are more susceptible to damage or degeneration. Multiple recent studies suggest that glutamatergic neurons in L2/3 of mouse visual cortex are particularly responsive to neuronal activity at the transcriptomic level^19,46^, thus the sensitivity of these cells to increased activity may be conserved between the two species.

### Blunted SST-mediated inhibition of IT-projecting neurons within the eME

While glutamatergic neurons underwent the strongest transcriptomic shifts within the eME, most GABAergic populations exhibited DEGs as well. Among GABAergic neurons, SST+ interneurons responded the strongest to the eME, upregulating 149 genes in focal compared to non-focal tissue (**Fig. 2a**). GO analysis revealed the induction of gene programs including IEGs (e.g. *EGR1* and *JUN*), stress- and synapse-associated factors such as heat-shock proteins (HSPs; e.g. *HSP90AA1* and *HSPA1A*), and apoptosis-associated genes (e.g. *BTG2*, *PRNP*, *MCL1*, and *CLU*) within these cells. Concomitantly, transcripts associated with development and cellular remodeling, which were enriched in excitatory neurons, were not induced in focally localized GABAergic cells, leading us to hypothesize that a loss of cellular fitness of SST+ GABAergic neurons may contribute to the dampening of inhibition within the eME (**Fig. 2f**). To further test this possibility, we harnessed the computational package *CellChat* which estimates the number and strength of putative cell-cell interactions in scRNAseq data based upon known ligand-receptor relationships^47^. Consistent with our hypothesis, *CellChat* revealed a decrease in the predicted strength of SST+ interactions with Exc_L2/3IT, Exc_L5IT_GRIN3A+, and Exc_L6IT glutamatergic neurons within the eME, suggesting that decreased SST-mediated inhibition may contribute to the strong transcriptomic sensitivity of IT-projecting glutamatergic cells to the eME (**Fig. 2g**).

To gain molecular insights, we asked which ligand-receptor pairs underlie these predicted changes in inhibition. We found that the attenuation of SST-to-IT projecting neuron communication was primarily driven by a dampening of FLRT2-mediated cell adhesion (**Fig. 2h**). As FLRT2 plays essential roles in neuronal migration and axon guidance^48^, the loss of this signaling module could reflect a role for dysregulated inhibition in activity-related (and/or disease-related) changes to intratelencephalic circuits. Together, these data suggest that GABAergic-to-glutamatergic neuron signaling is dampened within the eME, and that SST+ interneurons, the strongest GABAergic responders, contribute to this predicted loss of inhibition.

### Acute electrical stimulation triggers robust shared and cell-type-specific gene programs in human cortex

Presurgical and intraoperative mapping showed that the eME is characterized by high levels of seizure-associated activity compared to non-focal regions in our study’s patients, a feature consistent with the broad induction of canonical IEGs that we observe across numerous glutamatergic and GABAergic neuronal cell types. We speculate that the transcripts upregulated by cells in the eME may fall into two non-mutually-exclusive classes: (1) transcripts that are altered due to neuronal dysfunction and therefore represent disease signatures (and may be the most promising targets for therapeutic intervention); and (2) transcripts that are induced as part of a neuron’s normal response to heightened activity. For instance, the high levels of multi-functional IEGs such as *FOS* and *EGR1* that we observe in focal tissue may reflect a disease-associated response of the neuron to cellular stress and localized neuroinflammation, and/or might be a component of the neuron’s normal transcriptional response to high levels of synaptic activity. To begin to disambiguate activity-dependent transcription from disease-associated gene programs, we sought to understand how non-epileptogenic human brain cells respond to biphasic electrical stimulation *in vivo* (**Fig. 1a**). Although this stimulation paradigm falls short of recapitulating endogenous activity patterns in human cortex, we expect the gene signatures induced in neurons following electrical stimulation to at least partially overlap with gene programs organized by more physiologically relevant patterns of neuronal activity. Thus, we next performed differential gene expression analysis to identify transcriptomic differences between stimulated and unstimulated non-focal regions. We identified broad transcriptomic changes across numerous cell types, with 1,196 unique genes upregulated and 626 genes downregulated by stimulation in at least one cell type (**Fig. 3a**). Notably, these numbers are on par with the abundance of transcripts that are dysregulated within the eME compared to non-focal tissue, where we observed 1,301 and 711 genes up- or downregulated, respectively. As in our comparison of focal and non-focal regions, Exc_L2/3IT, Exc_L5IT_GRIN3A+, and Exc_L6IT neurons again exhibited the highest numbers of DEGs following acute stimulation even when cell numbers were downsampled to 500 cells per type (**Extended data Fig. 3b**), indicating that some of the transcriptomic changes observed in these cells within the eME may represent a normal response to activity rather than a vulnerability to disease (**Fig. 3a**). Among these, Exc_L2/3IT neurons were the top DEG-bearing subtype (**Fig. 3b**). Again, IT-projecting glutamatergic neurons induced a shared signature of IEGs, including *ARC*, *EGR1*, and *JUND* following acute stimulation (**Fig. 3c**). IEG induction following stimulation extended beyond IT-projecting neurons to other glutamatergic and GABAergic cell types, however the IEG response included fewer canonical IEGs and extended to fewer cell types than what was observed in the eME (**Fig. 3d,e**). Interestingly, we also observed the induction of *EGR1* in both microglia and oligodendrocyte precursor cells following acute stimulation but not within the eME (**Fig. 3d,e**), suggesting that glia are more responsive to acute stimulation than to epileptogenic activity. Glutamatergic, GABAergic, and glial cells robustly upregulate several HSPs (e.g. *HSPA1A*, *HSPA1B*, *HSPA8*, *HSP90AA1*, *HSP90AB1*, and *HSPH1*) following acute stimulation, a pattern observed in the eME as well (**Fig. 3f,g**). In this context, HSP induction could reflect a cellular response to stress, or may be more related to the initiation of activity-induced synaptic remodeling and neuroprotection, processes to which HSPs are known to contribute^49,50^.

**Figure 3.**
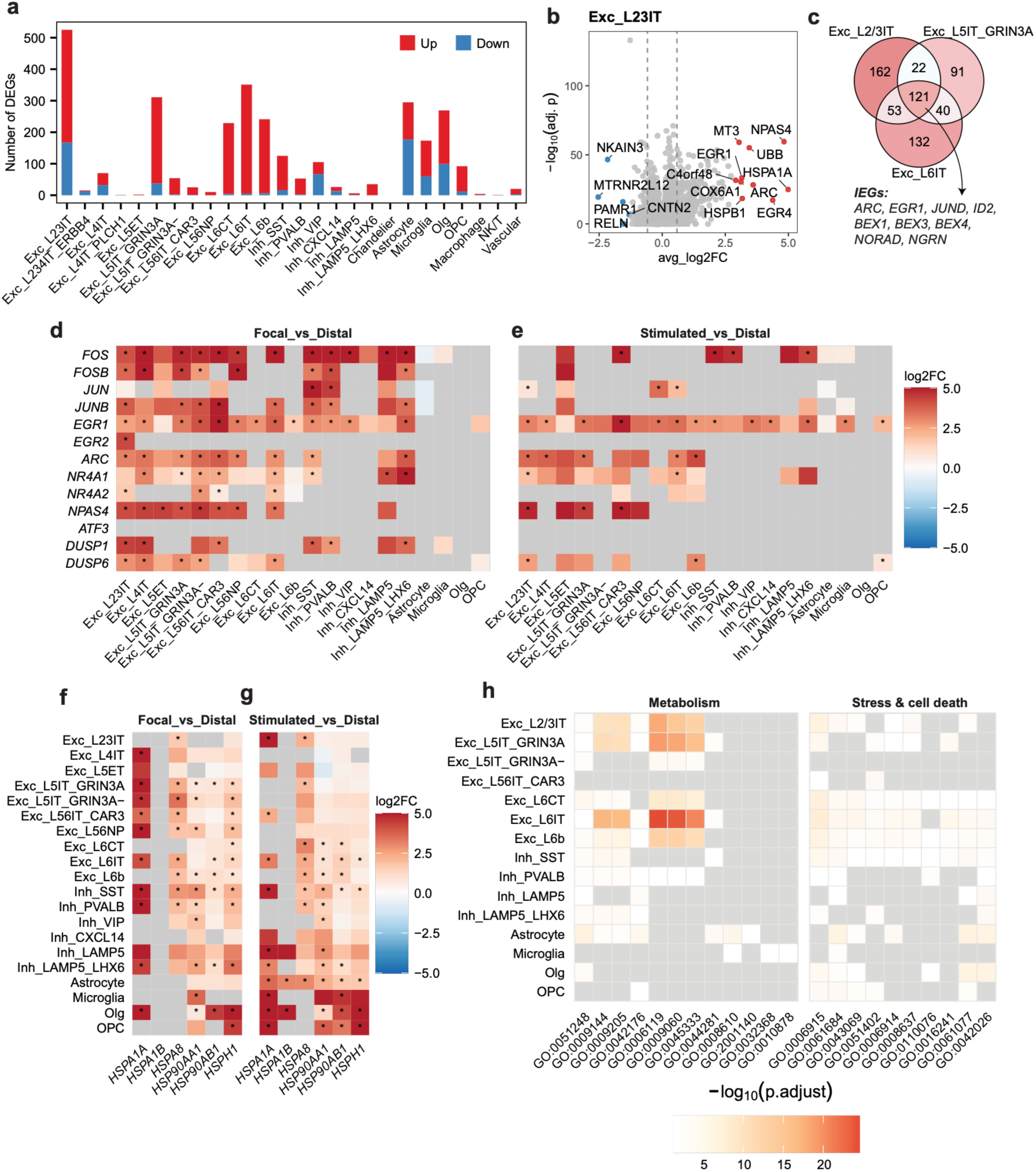
Acute electrode stimulation triggers shared and cell-type-specific gene programs in neuronal and glial cells. (a) Abundance of differentially expressed genes (DEGs) between stimulated and unstimulated non-focal regions, thresholded at FDR < 0.05 and |log2FC| > log2(1.5). Red, upregulated by stimulation; blue, downregulated by stimulation. (b) Volcano plot of DEGs in Exc_L2/3IT neurons between stimulated and non-focal regions. The top 10 DEGs upregulated by stimulation and the top 5 DEGs downregulated by stimulation are highlighted in red and blue, respectively, with all other genes shown in grey. Dashed lines indicate |log2FC| = log2(1.5) thresholds. (c) Venn diagram of stimulation-induced DEGs shared among Exc_L2/3IT, Exc_L5IT_GRIN3A, and Exc_L6IT neurons. Nine shared IEGs are highlighted. (d) Heatmap displaying differential expression of canonical IEGs and HSPs in focal compared to non-focal regions. Colors indicate log2FC as shown in the scale on the right, with warmer colors reflecting higher expression following stimulation. Asterisks denote significant genes with |log2FC| > log2(1.5) and FDR < 0.05. Non-significant genes are shown in gray; only genes with min.pct ≥ 0.1 were tested. (e) Same as (d) except IEGs induced by acute stimulation are plotted. (f) Heatmap displaying differential expression of HSPs in focal compared to non-focal regions. Warmer colors reflect higher expression following stimulation. Colors indicate log2FC as shown in the scale on the right, with warmer colors reflecting higher expression following stimulation. Asterisks denote significant genes with |log2FC| > log2(1.5) and FDR < 0.05. Non-significant genes are shown in gray; only genes with min.pct ≥ 0.1 were tested. (g) Same as (f) except HSPs induced by acute stimulation are plotted. (h) Heatmap of Gene Ontology (GO) enrichment across major cell types based on DEGs between stimulated and unstimulated non-focal regions. Significantly enriched GO terms (FDR < 0.05) are colored by −log10(FDR), with non-significant terms shown in grey.

While IEGs and HSPs were upregulated in both the eME and following acute stimulation, a unique feature of acute stimulation was the induction of robust gene programs in non-IT-projecting glutamatergic neurons residing in layer 6 (L6). Whereas non-IT projecting neurons in L6 (i.e. Exc_L6CT and Exc_L6b) upregulated 18 and 89 transcripts in focal compared to non-focal tissue, they upregulated 225 and 235 transcripts, respectively, following acute stimulation, with these transcripts revolving around stress-associated and proteostatic gene programs (**Fig. 3a and Extended Data Fig. 4**). Thus, at a population level, while IT-projecting glutamatergic neurons were the strongest responders to both the eME and acute stimulation, L6 glutamatergic neurons and glia exhibited preferential sensitivity to acute stimulation. As discussed further below, beyond IEGs, transcripts induced by acute stimulation in numerous cell types were associated with cellular stress pathways and metabolic mechanisms that support oxidative phosphorylation (**Fig. 3h**).

### Refining our understanding of disease signatures based on activity-dependent gene programs in non-focal cells

To begin to disambiguate disease and activity-regulated signatures, we next asked whether the transcripts that are acutely induced in major populations of glutamatergic and GABAergic neurons by acute stimulation overlapped with the transcripts that are upregulated within the eME. Across major glutamatergic and GABAergic neuronal classes, including IT neurons, L6 neurons, and representative GABAergic subtypes such as SST^+^ interneurons, each population displayed a subset of upregulated genes shared between acute and epileptogenic conditions. These overlapping transcripts prominently included IEGs such as *EGR1* and *NPAS4*, as expected. Notably, *FTH1* which encodes Ferritin Heavy Chain 1, an iron storage protein, was also among the most strongly upregulated genes across multiple cell types, including Exc_L5IT_GRIN3A, Exc_L6CT, Inh_PVALB, and Inh_SST neurons, suggesting that both chronic epileptogenic activity and acute stimulation converge on a transcriptional program that enhances neuronal iron handling and may increase vulnerability to ferroptotic stress (**Fig. 4a-d**).

**Figure 4.**
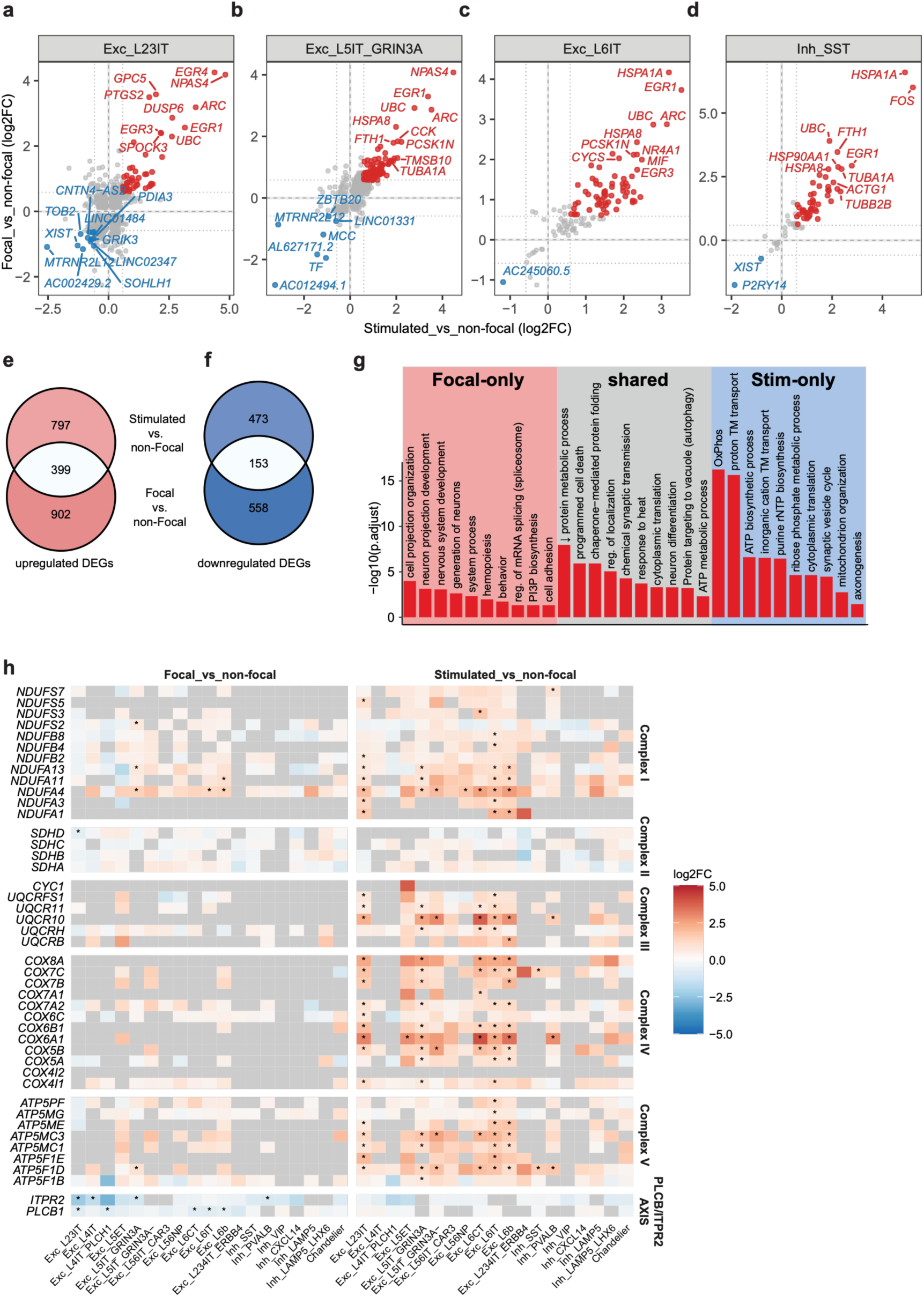
Disambiguating disease- and activity-regulated transcripts in the human brain. (a)-(d) Scatter plots of DEGs in Exc_L23IT (a), Exc_L5IT_GRIN3A (b), Exc_L6IT (c), and Inh_SST (d) neurons induced by both the eME and acute stimulation, as determined by MAST (Model-based Analysis of Single-cell Transcriptomics) using a generalized linear hurdle model, with patient identity included as a latent variable to control for inter-individual effects. The top 10 shared upregulated and top10 shared downregulated DEGs in each cell type are labeled with gene symbol. Upregulated and downregulated DEGs are highlighted in red and blue (|log2FC| > log2(1.5), FDR < 0.05), respectively. (e) Venn diagram displaying overlap between the 1,301 unique transcripts upregulated within the eME and the transcripts induced by acute stimulation. (f) Venn diagram displaying overlap between the 711 unique transcripts downregulated within the eME and the transcripts downregulated by acute stimulation. (g) Gene ontology (GO) analysis of upregulated shared and unique transcripts as plotted in (e). (h) Heatmap showing differential expression of mitochondrial electron transport chain (ETC) complex genes and PLCβ–ITPR2 axis components across major neuronal and non-neuronal cell types. The left panel shows focal versus non-focal regions, and the right panel shows stimulated versus non-focal regions. Genes are grouped by ETC complexes (Complex I–V) and the PLCβ–ITPR2 axis as indicated. Colors represent log2 fold change (log2FC) according to the scale on the right, where warmer colors indicate higher expression relative to the non-focal region. Asterisks denote significantly differentially expressed genes with |log2FC| > log2(1.5) and FDR < 0.05, as determined by MAST (Model-based Analysis of Single-cell Transcriptomics) using a generalized linear hurdle model, with patient identity included as a latent variable to control for inter-individual effects. Non-significant genes are shown in gray; only genes expressed in at least 10% of cells in either group (min.pct = 0.1) were tested.

To investigate the relationship between eME-enriched and acutely activated gene programs more systematically beyond this subset of cell types, we next quantified overlap between the 1,301 unique transcripts upregulated in at least one cell type within the eME and the 1,196 transcripts upregulated in at least one cell type by acute stimulation. Strikingly, we found that about 31% of the transcripts upregulated within the focal compared to the non-focal region were also induced following acute stimulation (**Fig. 4e**). This substantial degree of overlap suggests that some transcripts that are more highly expressed within the eME are upregulated in response to heightened activity, not necessarily as a result of the tissue’s pathological state. Conversely, about 22% of the genes downregulated in the eME were also downregulated following acute stimulation, indicating greater overlap between upregulated than between downregulated transcripts (**Fig. 4f**). Finally, when we assessed the overlap between eME and stimulation-dependent gene programs on a cell-type-specific basis, we found that gene programs induced in GABAergic neurons were more likely to be shared than those induced in glutamatergic neurons (**Extended data Fig. 5**).

To shed light on the nature of the transcripts that are upregulated by both epileptogenic and acute activity, we performed GO analysis on the cohort of shared genes specifically. We found that these transcripts were enriched for functional processes such as chaperone-mediated protein folding, chemical synaptic transmission, and cytoplasmic translation, all processes known to be orchestrated by neuronal activity (**Fig. 4g**). Conversely, the transcripts that were upregulated within the eME but unchanged following acute stimulation were enriched for functions such as neuron projection development, nervous system development, mRNA splicing, and cell adhesion. These results suggest that both the eME and acute stimulation promote the expression of transcripts with defined roles in activity-dependent processes, while the eME selectively initiated a gene program associated with neuronal development. Finally, as discussed further below, the transcripts that were upregulated by acute stimulation but unchanged by the eME were enriched for functions such as oxidative phosphorylation, ATP biosynthesis, cytoplasmic translation, and synaptic vesicle cycling (**Fig. 4g**). In summary, about 1/3 of the transcripts that are enriched within the eME are also upregulated by acute stimulation, suggesting that a fraction of epilepsy-associated gene expression reflects conserved responses to heightened activity rather than, or in addition to, disease-specific programs.

### Glutamatergic neurons within epileptogenic cortex fail to metabolically adapt to the energy demands of neuronal activation

While transcripts that are induced in the eME but not following acute stimulation are more likely to represent disease signatures that evolve over time than activity-regulated factors, we reason that transcripts that are induced by stimulation but unchanged by epileptogenic activity likely fall within one of two categories: (1) transcripts that are rapidly but *transiently* induced by neuronal activity, and (2) transcripts that are normally induced by activity but fail to be induced in the eME, potentially due to disease. For example, if healthy neurons adapt to neuronal stimulation by upregulating factors such as synaptic remodelers, a lack of this response in epileptogenic-localized neurons may reflect a disease-associated inability to properly adapt. In our dataset, as mentioned briefly above, we identified a module related to ATP production that clearly exhibits the second pattern.

Neuronal activity is of high energetic demand, especially in the human brain^51,52^, and research shows that oxidative phosphorylation in the mitochondrion is one major energy source underpinning this activity^53^. Consistent with synaptic innervation rapidly engaging metabolic pathways to support this function, we observed a coordinated induction of electron transport chain (ETC) proteins across glutamatergic neurons following acute stimulation. The ETC is a family of five protein complexes residing in the inner mitochondrial membrane that transfer electrons from NADH and FADH_2_ to oxygen, generating an electrochemical gradient that produces ATP. We observed a strong, coordinated induction of several ETC components within each complex (interestingly, except for complex II) in response to acute biphasic stimulation. In contrast, ETC expression was largely normal in focal compared to non-focal tissue (**Fig. 4h**). This observation suggests that neurons within healthy cortical tissue rapidly upregulate ETC expression within thirty minutes of stimulation to increase ATP production thereby fueling their functions, while neurons within epileptogenic tissues, despite inducing high levels of activity-regulated IEGs, are unable to metabolically adapt.

What upstream factors might contribute to the inability of neurons within epileptogenic cortex to increase ETC expression? Recent work suggests that a PLCβ–IP₃–IP₃R pathway may play a role^54,55^. Upon activation by Gq proteins, cytosolic PLCβ produces the second messengers DAG and IP3. IP3 binds the ITPR2 calcium channel on the endoplasmic reticulum near mitochondria, driving calcium into the mitochondrial membrane to increase ATP production. Notably, we observed a selective downregulation of *PLC*β*1* in Exc_L23IT, Exc_L4IT_PLCH1, Exc_L6CT, Exc_L6IT, and Exc_L6b neurons in focal compared with non-focal regions. In parallel, *ITPR2* was downregulated in Exc_L23IT, Exc_L4IT, Exc_L5IT_GRIN3A, and Inh_PVALB neurons, further indicating suppression of the PLCβ–IP₃–IP₃R axis as a potential pathway underlying the attenuated ETC gene program in focal neurons. Notably, *PLC*β*1* and ITPR2 expression were unchanged by acute stimulation, further suggesting that dampened activity of this pathway may contribute to the inability of neurons within the epileptogenic focus to respond (**Fig. 4h**).

### Reactive astrocytes blunt inflammatory signaling while microglia exhibit signs of activation within the eME

In addition to glutamatergic and GABAergic neurons, brain-resident glial cells also contribute significantly to cortical function and have been implicated in DRE^56^. Our data show that, after IT-projecting glutamatergic neurons, astrocytes are the fourth most strongly affected cell type in both epileptogenic and acute conditions (**Fig. 2a and 3a**). To interrogate astrocytic changes in greater detail, we re-clustered all astrocytes in the dataset to reveal three distinct subtypes: homeostatic astrocytes, reactive astrocytes (RAs, characterized by high levels of *CD44*), and lipid-accumulating reactive astrocytes (LARAs, characterized by high levels of *APOE*; **Fig. 5a**). Scoring against previously reported RA and LARA signatures yielded highly reproducible classification of these states^57–61^. Compared to homeostatic astrocytes, we find that LARAs express higher levels of transcripts involved in oxidative phosphorylation, aerobic respiration, mitochondrial ATP synthesis, cytoplasmic translation, and ion transport. Conversely, RAs express higher levels of transcripts involved in cell adhesion (eg *CD44*, *NRXN3*) and cytokine and immune responses (e.g. *TLR4*, *STAT1*, *CCL2)*(**Fig. 5b,c**). Interestingly, despite expressing higher levels of inflammatory transcripts than their homeostatic counterparts, RAs have been previously reported to have some neuroprotective functions^58^. Our data are consistent with RAs playing a protective role within the eME, as GO analysis revealed that RAs downregulated transcripts related to cytokine signaling and other inflammation-related functional categories within focal compared to non-focal tissue (**Fig. 5d**).

**Figure 5.**
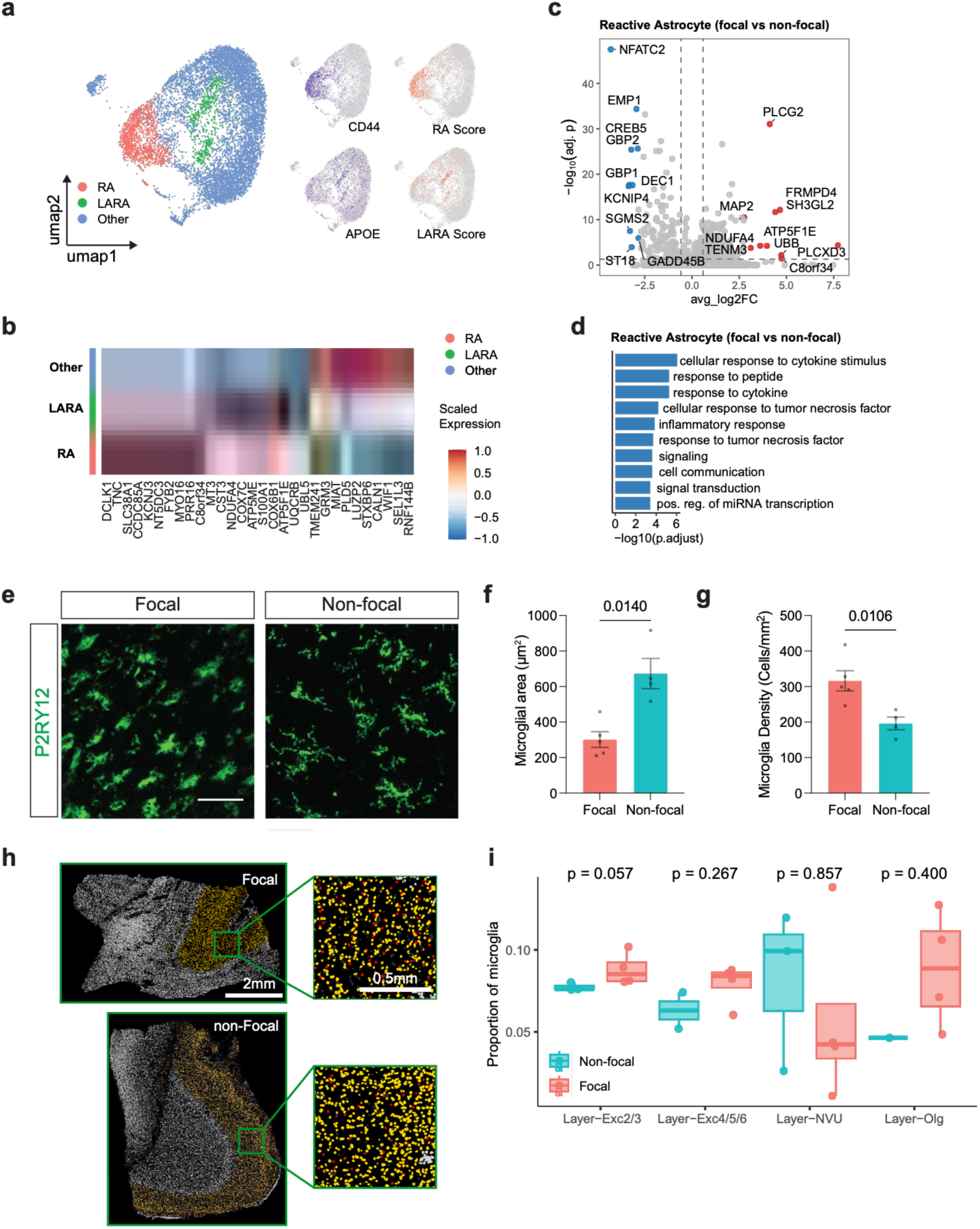
Reactive astrocytes and microglia balance neuroinflammation within epileptogenic cortex. (a) Uniform Manifold Approximation and Projection (UMAP) plot of astrocytes following hierarchical sub-clustering, colored by subtypes (left). Feature plots showing expression of *CD44* and *APOE* corresponding with RA and LARA scores (right). (b) Heatmap displaying the top 10 most highly enriched genes within each astrocytic subtype, colors indicate gene-wise z-score–normalized expression across astrocytic subtypes. (c) Volcano plot of DEGs in reactive astrocytes between focal and non-focal regions. The top 10 DEGs upregulated in focal and the top 10 DEGs downregulated in focal are highlighted in red and blue, respectively, with all other genes shown in grey. Dashed lines indicate |log2FC| = log2(1.5) thresholds. (d) Gene ontology terms enriched in reactive astrocytes in focal regions compared with non-focal regions. The y axis indicates −log10(FDR). (e) Confocal images of cortical sections of focal and non-focal regions immunostained for the microglial marker IBA1 (green). Scale bar, 100 μm. (f) Average area of individual microglia in focal versus non-focal regions. (g) Average density of microglia in focal versus non-focal regions. (f),(g) unpaired t test, n = 5 focal and 4 non-focal regions. (h) Example images of Xenium data showing microglia (red) within the L2/3 niche. Scale bar, 0.5 mm. (i) Box plots of microglial proportions across cortical layers in Xenium data in focal versus non-focal cortex. Each point represents one sample. Boxes indicate median and interquartile range, with whiskers extending to 1.5× IQR. P values were calculated using a two-sided Wilcoxon rank-sum test within each layer. Red, focal; blue, non-focal.

Astrocyte reactivity is promoted by signals from microglia^59^, leading us to next assess microglial responses to epileptogenic and acute activity. In response to acute stimulation, microglia upregulate robust cohorts of genes involved in inflammatory signaling and chemotaxis (e.g. *IL1B*, *CCL3*, *CCL4*, *ADGRE2*, and *FLT1*), synaptic organization and axon guidance (e.g. *HOMER1*, *DLGAP4*, *PTPRD*, and *PLXNA2*), and neurodegeneration (e.g. *SPP1*, *LPL*, *PPARG*, *MSR1*, and *ABCB4*). Although microglia did not exhibit a large number of DEGs in focal compared to non-focal regions, immunofluorescence revealed that they assume an amoeboid morphology (classically associated with immune activation in macrophages) and increase their density within the eME (**Fig. 5e-g**). Furthermore, spatial analysis via the Xenium platform largely corroborated the increased density of microglia within the focal region, especially within the layer 2/3 compartment (**Fig. 5h,i**). Together, these findings indicate that microglia shift from a homeostatic toward a cytotoxic state within the eME, but retain healthy properties (i.e. a highly ramified morphology, lower density) in anatomically matched regions exhibiting non-epileptogenic levels of activity. In parallel, RAs within the eME decreased inflammatory signaling suggesting a potential neuroprotective role.

### Circulating CD14+ monocytes exhibit heightened activation in individuals with DRE

Consistent with a growing number of studies revealing a strong neuroinflammatory component to epilepsy, our data show that microglia exhibit signatures of inflammation that are localized within the eME. As microglia are peripherally derived macrophages, this led us to wonder whether inflammatory signatures of DRE extend beyond the brain. To answer this question at single-cell resolution, we profiled peripheral blood mononuclear cells (PBMCs) from four DRE patients whose cortical tissue was included in the study, and seven healthy donors (**Fig. 6a**). Using pooled scRNA-seq with SNP-based genetic demultiplexing, we generated 189,110 high-quality cells spanning 25 immune and hematopoietic cell types to provide a comprehensive snapshot of the peripheral immune landscape in DRE (**Fig. 6b,c and Extended Data Fig. 6**).

**Figure 6.**
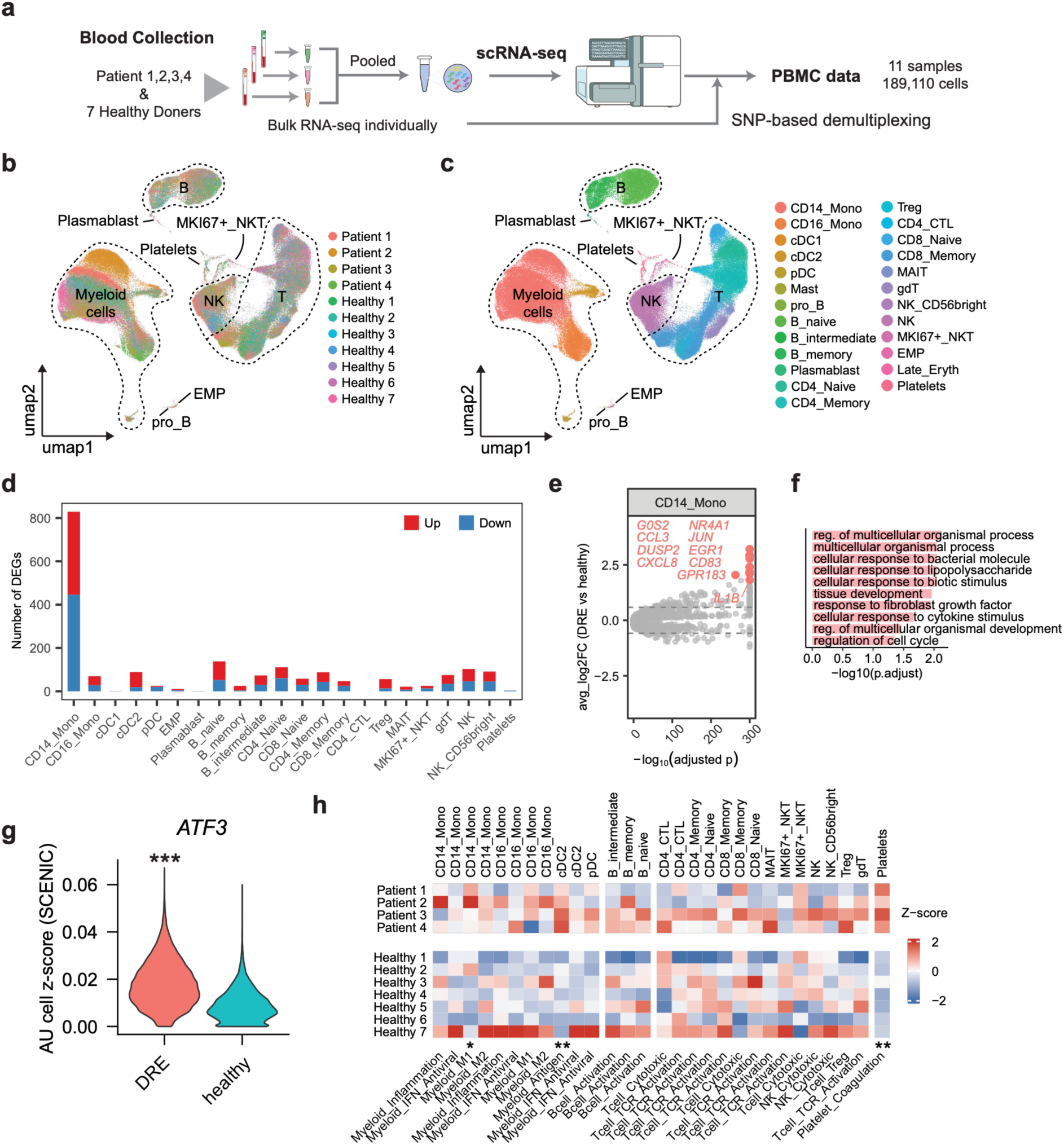
Circulating CD14+ monocytes display signs of activation in individuals with DRE. (a) Experimental approach to identify gene expression changes in peripheral blood mononuclear cells (PBMCs) isolated from four individuals in the study or seven healthy controls. (b) Uniform Manifold Approximation and Projection (UMAP) plot of all cells in the dataset colored by individual donor, legend in bottom right. (c) As in (b) but colored by cell type. (d) Abundance of differentially expressed genes (DEGs) between DRE and control individuals, thresholded at FDR < 0.05 and |log2FC| > log2(1.5). Red, upregulated in DRE; blue, downregulated in DRE. (e) Volcano plot of DEGs in CD14⁺ monocytes between DRE and healthy controls. The top 10 DEGs upregulated in DRE are highlighted in color, with all other genes shown in grey. Dashed lines indicate |log2FC| = log2(1.2) thresholds. (**f)** Top 10 GO terms enriched within DEGs of CD14⁺ monocytes upregulated in DRE versus healthy conditions. The y axis indicates −log10(FDR). (**g)** Violin plot of predicted ATF3 regulon activity in CD14⁺ monocytes from peripheral blood between DRE and healthy conditions, inferred by SCENIC. **p < 0.001 (two-sided Wilcoxon rank-sum test). (**h)** Heatmap of gene-set module scores across PBMC cell subtypes in DRE patients and healthy donors. Values are shown as column-wise Z-scores. Columns are grouped by major immune lineages (Myeloid, B cell, T/NK and Platelet) and labeled by the corresponding module. Asterisks indicate DRE–healthy differences for each subtype–module pair (two-sided Wilcoxon rank-sum test; *P < 0.05, **P < 0.01, ***P < 0.001).

We then performed cell type–resolved differential gene expression analysis between DRE and healthy individuals across all cell types, including myeloid cells, B cells, T cells, natural killer (NK) cells, and platelets. CD14⁺ monocytes exhibited by far the largest DEG set among all cell types, upregulating 383 while downregulating 446 transcripts in individuals with DRE compared to healthy controls (**Fig. 6d**). The DEGs in CD14+ monocytes which harbor the highest fold changes of upregulation in DRE individuals include IEGs (*JUN*, *EGR1*, *NR4A1*, *DUSP2*), pro-inflammatory and chemotactic factors (*IL1B*, *CCL3*, *CXCL8*), and markers of activation (*CD83*, *GPR183*; **Fig. 6e,f**). Single-cell regulatory network inference and clustering (SCENIC) revealed the transcription factor ATF3 as a potential mediator of these DRE-associated changes in circulating myeloid cells (**Fig. 6g**).

To more systematically predict the functional impact of transcriptomic changes across peripheral immune cells, we performed cell type–specific pathway scoring based on curated gene modules, with a focus on myeloid inflammation and polarization (M1/M2), antigen presentation, interferon response, lymphocyte activation, and platelet coagulation. CD14⁺ monocytes displayed elevated M1 polarization scores, consistent with a chronic pro-inflammatory state. Conventional dendritic cells (cDC2 cells) showed increased antigen-presentation activity, indicating enhanced potential for adaptive immune priming. Platelets demonstrated strong activation of coagulation pathways, suggesting a pro-thrombotic phenotype that may be associated with vascular inflammation and possible blood–brain barrier disruption. Other immune populations remained largely unchanged (**Fig. 6h**). Together, these data demonstrate that while the overall PBMC landscape is largely stable in DRE, specific subsets of cells undergo targeted activation, suggesting that peripheral immune perturbations may interact with vascular pathways and potentially influence brain–immune communication in the epileptic state.

## Discussion

This study introduces a single-cell transcriptomic atlas of epileptogenic and acute stimulation-dependent gene programs across 26 cell types in the human brain. In total, we identify 2,098 and 1,184 transcripts that were up- or downregulated by either epileptogenic activity or acute electrode stimulation in at least one cell type, respectively. We observe striking diversity in the responses of cells to either condition, such that some cells are virtually unchanged while others (e.g. IT-projecting glutamatergic neurons) undergo extensive transcriptomic remodeling. Among those that respond, the majority of transcripts impacted by activity were upregulated and cell-type-specific, except for shared IEG and HSP signatures. Notably, despite robust induction of activity-associated transcriptional programs, cells within the epileptogenic focus exhibited a decoupling between neuronal activity and mitochondrial energy responses related to oxidative phosphorylation. We additionally found that 31% of the transcripts that were enriched within the eME were also induced following acute stimulation, suggesting that the 69% of non-overlapping transcripts may be more likely to be disease signatures than a part of the cell’s normal response to increased activity (although functional experiments are needed to test this hypothesis). Expanding our analysis to glial and immune cells, we identify both local and systemic inflammatory signatures associated with DRE. Altogether, these findings advance our knowledge of activity-dependent transcription in the human brain, and provide mechanistic insights into the pathophysiology of neurological disorders that are caused by heightened activity.

While prior studies have revealed significant insights into DRE through scRNAseq^38–42^, our study is unique in several important ways. First, most studies analyzed paraffin-fixed brain samples collected by pathologists then subjected to long-term storage, which can lead to tissue degeneration and the introduction of confounding artifacts. Conversely, our analyses were performed on fresh brain tissue immediately flash frozen within minutes of resection. Second, other studies compare epileptogenic tissue from individuals with DRE to cortical tissue from healthy controls. Beyond the presence or absence of epileptogenic activity, this approach introduces unrelated experimental variables—e.g. sex, age at collection, age at disease onset—which could complicate interpretation. Our study eliminates these variables by comparing region-matched tissues with or without epileptogenic pathology harvested from the same individuals. Third, while other studies tend to focus on distinct cell types (e.g. neurons or immune cells), our unbiased analysis spans 26 cell types across neuronal and non-neuronal classifications. Most importantly, our study harnesses electrical stimulation of non-epileptogenic cortex prior to surgical resection (i.e. *in vivo* stimulation) to identify acute activity-dependent gene programs in the human brain, allowing us to more precisely pinpoint possible disease signatures. Thus, our study is particularly well-poised to define the eME through coordinated analysis of distinctly impacted brain regions harvested from unique individuals.

While these differences endow our study with novel insights, several caveats warrant consideration. For example, the number of patients included in our study is limited to six, and we were unable to collect all sample types from every patient due to clinical and surgical considerations. Thus, extending our findings to a larger cohort would likely expand and refine these insights. Second, DRE can be a profoundly debilitating disorder, and it is likely that even non-epileptogenic brain tissue from individuals with DRE will harbor some disease signatures, even though the activity signatures of epileptogenic regions were carefully mapped prior to and/or in the course of surgery. Thus, gene expression differences between epileptogenic and non-epileptogenic tissue may incompletely reflect differences between epileptogenic and healthy tissue, or may reveal false positives. Relatedly, while acute electrode stimulation clearly activates cortical cells as evidenced by wide-spread IEG induction, this mode of stimulation is likely harsher and differently patterned than endogenous neuronal activity. Thus, it is important to note that these analyses do not interrogate gene programs regulated by endogenous activity in the human brain, but rather genes that are impacted by chronic epileptogenic activity/pathology and robust, localized electrical stimulation. Finally, while the epileptogenic and non-epileptogenic samples were derived from tissue within the same resection cavity, there may still be subtle regional differences contributing to the transcriptomic changes that we report. Overall, despite these caveats, our study provides key insights into activity-dependent gene expression in the human brain and its relevance to epileptic pathology.

## Materials and Methods

### Human subjects

This study was approved by the Institutional Review Board at Northwell Health (IRB 20-0150), and all patients provided written informed consent prior to enrollment.

Patients with drug-resistant epilepsy (DRE), defined as failure to achieve sustained seizure control following adequate trials of two or more appropriately chosen antiseizure medications^62^ were recruited for this study. Prior to definitive surgical intervention, all patients underwent comprehensive presurgical evaluation including multimodal neuroimaging, scalp electroencephalography (EEG), and invasive intracranial EEG (iEEG) monitoring with depth and/or subdural strip electrodes, and extended monitoring for precise localization of the seizure onset zone. The decision to proceed with surgical resection was made through multidisciplinary epilepsy conference review and shared decision-making with each patient.

Six patients were enrolled in this study with an average age of 37.7 ± 12.2 years. The average duration of epilepsy prior to this surgical intervention was 19.3 ± 5.2 years.

| Subject ID | Age (years) | Sex | Seizure Onset (years) | Frequency of Seizures (n/month) | Prior Surgical History | Etiology | Location of Epileptogenic Foci |
| --- | --- | --- | --- | --- | --- | --- | --- |
| ID1 | 33 | Male | 13 | 5 | None | MTS | Left MTL |
| ID2 | 38 | Female | 10 | 2 | None | MTS | Left MTL |
| ID3 | 42 | Female | 25 | 1 | Right Parietal seizure focus resection; Right hippocampal laser interstitial ablation; Right temporal lobectomy | Post Hemorrhagic Stroke | Right Insula |
| ID4 | 59 | Female | 44 | 2 | Subtemporal amygdalohippocampectomy | MTS with additional foci in Insula | Left Insula |
| ID5 | 30 | Female | 8 | 2 | Vagal nerve stimulation | TLE | Left anterior Temporal lobe |
| ID6 | 24 | Male | 10 | 1 | Resection of ruptured frontal arteriovenous malformation | Post Hemorrhagic Stroke | Right Frontal lobe |

### Epilepsy surgery and human brain tissue collection

Etiology and surgical procedures were performed with standard neurosurgical techniques and tailored to each patient’s planned resection. Standard optical neuronavigation was used where necessary to localize epileptogenic foci and non-epileptogenic tissue as defined from presurgical workup. Non-epileptogenic tissue was designated for resection based on clinical and surgical indications. All tissue samples were flash frozen within ∼20 mins of resection.

### Intraoperative cortical stimulation

Acute intraoperative stimulation was performed only in brain tissue not suspected to be epileptiform. For example, in a scenario where the epileptogenic focus is deep (i.e. hippocampus or amygdala), surgical resection typically includes the anterior temporal lobe. Therefore, the cortical surface of the anterior temporal lobe designated is not suspected to be “epileptogenic” but surgically indicated as a part of the resection. Therefore, this portion of the brain was stimulated for the experiment.

All samples were stimulated immediately prior to resection using the previously described Penfield method^63^. A handheld bipolar ball tip probe (Ojemann Cortical Stimulator; Integra LifeSciences, Princeton, NJ) was used to apply focal biphasic stimulation with a maximum current of 10 mA, corresponding to a 20mA differential. Long-train bipolar stimulation was delivered at a frequency of 60 Hz with a pulse width of 500 μs and a train duration of 5s.

### Nuclei isolation for snRNA-seq

Brain tissue were transferred from a pre-chilled liquid nitrogen system to a 1 ml dounce homogenizer containing 300 µl of ice-cold supplemented lysis buffer (LB; 10mM Tris-HCl pH 7.4, 10mM NaCl, 3 mM MgCl2 6H2O, 0.05% IGEPAL NP-40, and 0.3U/uL protector RNAse inhibitor in ddH2O). The tissue was homogenized with a loose pestle then a tight pestle about 10-15 times each. The homogenate was then transferred to a 15mL falcon tube and was incubated on ice with an additional 3mL of LB. 5mL of Nuclear wash buffer (NWB; 5% BSA, 0.25% glycerol vol/vol, and 0.08U/uL Protector RNAse inhibitor in a 1:1 1X PBS:ddH2O solution) was added to the homogenate and gently inverted to mix. Homogenate was strained into a 40mL falcon tube with a 70um filter, centrifuged (5 min. at 500g (9/9 acc/dec) at 4°C), decanted, and resuspended with 10mL of NWB. Homogenate was then filtered with a 40um strain, centrifuged, decanted, and resuspended in 1mL of NWB. Homogenate was mixed with iodixanol to achieve a 50% iodixanol 2mL solution and gently transferred on top of a 2mL 29% optiprep cushion. Sample was centrifuged (30 min. at 4640 g (9/9 acc/dec) at 4°C), decanted and resuspended in final resuspension buffer (2.5% BSA, 0.25% glycerol vol/vol, and 0.3U/uL protector RNAse inhibitor in a 1:1 1X PBS:ddH2O solution) before proceeding with downstream sequencing.

### Xenium tissue section preparation

Brain samples were embedded in pre-chilled OCT with liquid nitrogen and Xenium slides were pre-chilled to -20°C in the cryostat chamber. OCT-embedded samples were sectioned on a Leica cryostat. A 10 μm-thick section was then mounted onto the imaging area of the Xenium slide using pre-cooled paintbrushes. Section was thaw-mounted by gently pressing finger underneath the slide where the tissue had been placed.

### PBMC isolation

PBMCs were isolated from human EDTA whole blood by density-gradient centrifugation using Ficoll (Ficoll-Paque™ PREMIUM, Cytiva; 1.078 g/mL).Whole blood (8 mL) was diluted to 50 mL in wash solution, layered onto Ficoll (14 mL per tube), and centrifuged at 400 x g for 30 min at 20–25°C (acceleration=1, deceleration=1, brake off). The PBMC-containing interphase was collected and washed three times in wash solution (320 x g, 10 min, ∼10°C; acceleration=8, deceleration=8), discarding the supernatant and resuspending the pellet between washes. For cryopreservation, cells were mixed 1:1 with freezing medium (CryoStor C55 Stemcell Technologies, 07933), frozen overnight at −80°C in a controlled-rate freezing container.

### Single-Nucleus RNA sequencing (snRNAseq)

Nuclei were stained with ViaStain AOPI (Nexcelom #CS2-0106-5mL) and counted using a Countess FL II automated cell counter. Single nucleus suspensions were loaded into a 10X Chromium X instrument targeting 10,000 recovered nuclei according to the manufacturer’s instructions. Barcoding and library preparation were performed using the NextGEM Single-Cell 3’ Library Kit v3.1(1000121; 10X Genomics). cDNA and libraries were checked for quality on Agilent TapeStation, and quantified by KAPA qPCR. Libraries were sequenced on a NextSeq2000 (Illumina) using the following read format: (28×10×10×90bp), to an average depth of approximately 25,000 reads per cell.

### Spatial Transcriptomics

Xenium *in situ* gene expression analysis was carried out using the human brain gene expression panel (10X Genomics #1000599) supplemented with a 100-gene custom add-on panel using Xenium instrument software version 3.0.2.0 and analysis software version 3.0.0.15. During panel compilation, a small subset of add-on targets conflicted with pre-allocated decoding codewords (including control-probe codewords) and was automatically flagged as *deprecated_codeword* entries; these targets were excluded from downstream analyses to prevent ambiguous transcript decoding. Across all embedded sections, the decoded transcript density ranged from 13 to 464 transcripts per 100 µm² across 11 regions.

### Single-cell RNA sequencing (scRNAseq)

Single-cell suspensions of PMBCs were washed and resuspended in PBS + 0.04% BSA, and an aliquot was stained with ViaStain AOPI (Nexcelom #CS2-0106-5mL) and counted using a Countess FL II automated cell counter. Single cell suspensions were loaded into a 10X Chromium X instrument targeting 20,000 recovered cells according to the manufacturer’s instructions. Barcoding and library preparation were performed using the GEM-X Universal Gene Expression v4 kit (1000691; 10X Genomics). cDNA and libraries were checked for quality on Agilent TapeStation, and quantified by KAPA qPCR. Libraries were sequenced on a NextSeq2000 (Illumina) using the following read format: (28×10×10×90bp), to an average depth of approximately 25,000 reads per cell.

### Data preprocessing

Raw sequencing data were processed using Cell Ranger (v7.0.0) with the GRCh38 reference genome (refdata-gex-GRCh38-2020-A). Ambient RNA contamination was removed using CellBender, and the corrected count matrices were imported into Seurat (v5) for downstream analysis.

#### Quality Control

For brain-derived nuclei, we retained cells with more than 500 detected genes, more than 1,000 total UMI counts, mitochondrial gene content below 2.5%, and a log10-transformed genes-per-UMI ratio greater than 0.8. Doublets were identified and removed using scDblFinder with default parameters. Only high-quality singlet cells passing all filtering criteria were retained for downstream analyses.

For blood-derived cells, we retained cells with more than 500 detected genes, total UMI counts between 500 and 20,000, mitochondrial gene content below 10%, ribosomal gene content below 50%, and a log10-transformed genes-per-UMI ratio greater than 0.8. To identify doublets, we performed genetic demultiplexing using vireo, which infers donor identity for each cell by comparing cell-specific SNP allele counts to donor genotype information. Cells assigned as doublets, indicating mixed genetic contributions from multiple donors, and cells labeled as unassigned were removed prior to downstream analyses.

#### Cell annotation

Cell types were annotated based on the expression of established canonical marker genes. For the brain dataset, neuronal populations were first classified into GABAergic/inhibitory and glutamatergic/excitatory neurons. GABAergic neuronal subtypes were identified using markers including *SST*, *PVALB*, *VIP*, *CXCL14*, *LAMP5*, and *LHX6*, distinguishing major interneuron classes. Glutamatergic neurons were annotated based on layer- and projection-associated markers, including *SLC17A7* as a pan-excitatory marker, CUX2 and RORB for upper-layer IT neurons, *PLCH1* for L4 IT neurons, and *BCL11B*, *FEZF2*, *GRIN3A*, *TLE4*, and *TSHZ2* for deep-layer L5–L6 excitatory neuron subtypes. Non-neuronal cell types were annotated using well-established markers, including PDGFRα for oligodendrocyte precursor cells, *MOG* for oligodendrocytes, *CX3CR1* for microglia, *CD163* for macrophages, *EBF1* for vascular-associated cells, and *PTPRC* for immune cells. Cell type identities were assigned based on the combined enrichment patterns of these marker genes, considering both expression level and the proportion of expressing cells across clusters.

Peripheral blood immune cell types were annotated based on the expression of established canonical marker genes. Major myeloid populations were identified using *CD14* and *FCGR3A* to distinguish classical and non-classical monocytes, respectively, while dendritic cell subsets were annotated based on *CLEC9A*, *FCER1A*, and *CLEC4C* expression. B cells were identified by *MS4A1*, *CD79A*, and *CD37*, with plasma cells annotated based on IGH genes. T cell populations were defined by *CD3D* expression and further subdivided into naïve and memory subsets using *LEF1*, *IL7R*, and *CCR7*. Regulatory T cells were identified by *FOXP3*, while cytotoxic T cell populations were annotated based on *CD8A*, *GZMB*, and *NKG7*. Natural killer (NK) cells were defined by *NKG7*, *KLRD1*, and *NCAM1* expression. Proliferating immune cells were identified by *MKI67*, and platelet populations were annotated based on *PPBP* expression.

#### Normalization, Batch effect correction, and DEG analysis

Gene expression counts were normalized using Seurat’s NormalizeData function, followed by identification of highly variable genes with FindVariableFeatures. The data were then scaled and centered using ScaleData, and principal component analysis (PCA) was performed on the scaled expression matrix.

To correct for batch effects and integrate data across samples, dimensionality reduction was harmonized using Harmony via the IntegrateLayers framework in Seurat (v5). Harmony integration was applied to the PCA embeddings, generating a corrected low-dimensional representation that was used for downstream analyses.

Differential gene expression analysis was performed using the FindMarkers function in Seurat (v5). For each cell type, pairwise comparisons were conducted between conditions using the MAST hurdle model. Genes were tested if they were expressed in at least 10% of cells in either group, and only genes with an absolute log2 fold change greater than log2(1.5) were considered significant. To account for inter-individual variability, patient identity was included as a latent variable in the model. Resulting p values were adjusted for multiple testing using the Benjamini–Hochberg procedure.

### Xenium data analysis

10x Genomics Xenium output files were imported into Seurat (v5). Cell centroid coordinates and segmentation boundaries were obtained from the Xenium-provided cell annotation files, and molecule-level transcript coordinates were imported from the Xenium transcript parquet files. Only molecules with a quality value (QV) ≥ 20 were retained. Spatial information was assembled into Seurat field-of-view (FOV) objects for downstream spatial analyses.

Cell type annotation for Xenium-segmented cells was performed using RCTD (Robust Cell Type Decomposition; spacexr). The human brain snRNA-seq dataset generated as part of this project and processed using Seurat (v5) was used as the annotation reference.

To characterize local cellular neighborhoods, a niche assay was constructed for each Xenium region using Seurat’s BuildNicheAssay, grouping cells by the RCTD-predicted cell type. Neighborhood graphs were built using a fixed number of spatial nearest neighbors (neighbors.k = 30), and niches were defined by clustering local neighborhood compositions (niches.k as specified per region). For downstream layer-level analyses, predicted fine cell types were collapsed into coarse categories (e.g., inhibitory neurons, upper-layer excitatory neurons, deep-layer excitatory neurons, glia, and other immune/vascular populations), and niche calling was repeated using the coarse labels when appropriate.

For each region, niches were manually mapped to interpretable spatial layers based on the enrichment patterns of coarse cell type compositions (e.g., excitatory upper-layer, excitatory deep-layer, neurovascular unit, oligodendrocyte-enriched niche), generating a layer metadata field for visualization and comparative analyses.

### Immunofluorescence (IF)

Immunofluorescence staining was performed on flash-frozen human brain tissue sections using a protocol optimized for human cortical samples. Tissue was embedded in OCT immediately after resection and cryosectioned at 10 µm thickness. Sections were stored at −80 °C until use. Slides were post-fixed in 4% paraformaldehyde in 1× PBS for 30 min at room temperature, briefly washed in PBS, and treated with potassium permanganate working solution for 10 min to reduce autofluorescence. Sections were washed in PBS containing 0.1% Triton X-100 and incubated with BLOXALL blocking solution for 10 min to quench endogenous peroxidase activity, followed by additional washes.

Sections were blocked for 1 h at room temperature in blocking buffer consisting of 2.5% normal horse serum with 0.5% Triton X-100. Primary antibody solution was applied and sections were incubated overnight at 4 °C in a humidified chamber. The following primary antibody was used: rabbit anti-P2RY12 (Sigma-Aldrich, HPA014518), a microglia-specific marker validated for human tissue.

After primary incubation, sections were washed three times in PBS and incubated with Alexa Fluor 488–conjugated anti-rabbit IgG secondary antibody for 30 min at room temperature. Sections were washed again in PBS and nuclei were counterstained with DAPI. Coverslips were mounted using antifade mounting medium and stored at 4 °C until imaging.

### Statistics and reproducibility

No statistical methods were used to predetermine sample size. Sample sizes were based on the availability of surgically resected human tissue from patients with drug-resistant epilepsy. No data was excluded unless they failed predefined quality control criteria. Investigators were blinded to sample identity during data collection but not necessarily during analysis.

For single-nucleus and single-cell RNA-seq analyses, statistical tests were performed as described in the corresponding sections. Differential gene expression analyses were conducted using the MAST hurdle model implemented in Seurat (v5), with patient identity included as a latent variable to account for inter-individual variability. P values were adjusted for multiple comparisons using the Benjamini–Hochberg method.

For brain-derived datasets, only genes expressed in at least 10% of cells in either group and with an absolute log2 fold change greater than log2(1.5) were considered. For PBMC single-cell datasets, genes expressed in at least 10% of cells in either group with an absolute log2 fold change greater than log2(1.2) were considered.

For spatial transcriptomics analyses, statistical comparisons of cell-type proportions, niche composition, and spatial enrichment were performed using non-parametric tests unless otherwise specified in the figure legends.

All statistical analyses were performed using R (version 4.4.3). Exact statistical tests, sample sizes, and definitions of center and dispersion are indicated in the corresponding figure legends.

## Data Availability

All transcriptomic data and associated code are in the process of deposition to an appropriate repository that meets the constraints of the tissue collection as determined by the IRB consent form, which requires controlled access to our patient data. In the meantime we are happy to share any and all raw and processed data with the reviewers.

## Acknowledgements

We thank the following individuals for providing critical feedback on the manuscript: Dr. Michael Greenberg (Harvard), Dr. Ava Carter (Harvard), Dr. Derek Southwell (Duke), Dr. Marty Yang (UCLA), and Dr. Austin Ferro (CSHL). In addition we thank Drs. Xiaonan (Richard) Sun, David Bonda, and Prashin Unadkat with help in obtaining tissue samples. This work was performed in concert with the Single-Cell Biology and Next-Generation Genomics Shared Resource Facilities at Cold Spring Harbor Laboratory, which are supported by National Institutes of Health Cancer Center support grant 5P30CA045508. This work was supported by a Rita Allen Scholar Award, McKnight Scholar Award, Klingenstein-Simons Fellowship in Neuroscience, and a Brain and Behavior Foundation NARSAD grant (to L.C.).

## Notes

### Competing Interest Statement

The authors have declared no competing interest.

