## Extended Data for "Deciphering epileptogenic and activity-dependent gene programs in the human brain"

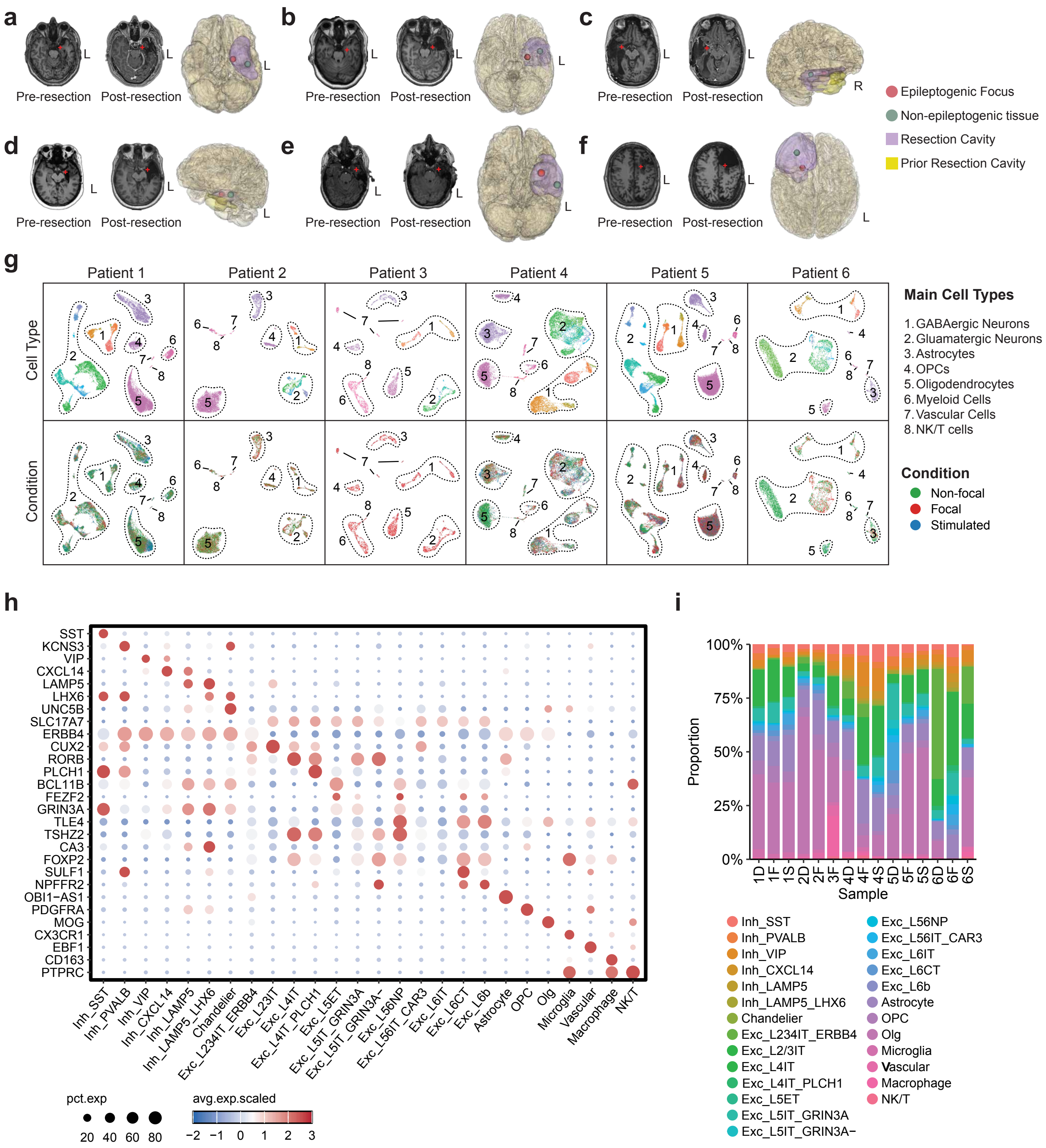

a
b
c
d
e
f
Pre-resection
Post-resection
L
R
Epileptogenic Focus
Non-epileptogenic tissue
Resection Cavity
Prior Resection Cavity
g
Patient 1
Patient 2
Patient 3
Patient 4
Patient 5
Patient 6
Cell Type
Condition
Main Cell Types
1. GABAergic Neurons
2. Gluamatergic Neurons
3. Astrocytes
4. OPCs
5. Oligodendrocytes
6. Myeloid Cells
7. Vascular Cells
8. NK/T cells
Condition
Non-focal
Focal
Stimulated
h
SST
KCNS3
VIP
CXCL14
LAMP5
LHX6
UNC5B
SLC17A7
ERBB4
CUX2
RORB
PLCH1
BCL11B
FEZF2
GRIN3A
TLE4
TSHZ2
CA3
FOXP2
SULF1
NPFFR2
OBI1−AS1
PDGFRA
MOG
CX3CR1
EBF1
CD163
PTPRC
Inh_SST
Inh_PVALB
Inh_VIP
Inh_CXCL14
Inh_LAMP5
Inh_LAMP5_LHX6
Chandelier
Exc_L234IT_ERBB4
Exc_L23IT
Exc_L4IT
Exc_L4IT_PLCH1
Exc_L5ET
Exc_L5IT_GRIN3A
Exc_L5IT_GRIN3A−
Exc_L56NP
Exc_L56IT_CAR3
Exc_L6IT
Exc_L6CT
Exc_L6b
Astrocyte
OPC
Olg
Microglia
Vascular
Macrophage
NK/T
pct.exp
20 40 60 80
avg.exp.scaled
−2 −1 0 1 2 3
i
100%
75%
50%
25%
0%
Proportion
1D 1F 1S 2D 2F 3F 4D 4F 4S 5D 5F 5S 6D 6F 6S
Sample
Inh_SST
Inh_PVALB
Inh_VIP
Inh_CXCL14
Inh_LAMP5
Inh_LAMP5_LHX6
Chandelier
Exc_L234IT_ERBB4
Exc_L2/3IT
Exc_L4IT
Exc_L4IT_PLCH1
Exc_L5ET
Exc_L5IT_GRIN3A
Exc_L5IT_GRIN3A−
Exc_L56NP
Exc_L56IT_CAR3
Exc_L6IT
Exc_L6CT
Exc_L6b
Astrocyte
OPC
Olg
Microglia
Vascular
Macrophage
NK/T

**Extended Data Figure 1. Anatomical visualization of resected brain regions and transcriptomic cell-type annotation across patients and conditions. (a–f)** Pre- and post-resection clinical imaging of tissue collected from six individuals with drug-resistant epilepsy (DRE), illustrating the epileptogenic focus and non-focal collection sites, color panel on right. **(g)** Uniform Manifold Approximation and Projection (UMAP) plots by cell type (top) and condition (bottom) across all six patients. Labels defined on the right. **(h)** Dot plot showing expression of canonical marker genes across annotated cell types. Dot size indicates the percentage of cells expressing each transcript, and color represents scaled average expression. **(i)** Stacked bar plots showing the proportion of major cell classes across samples, grouped by condition.

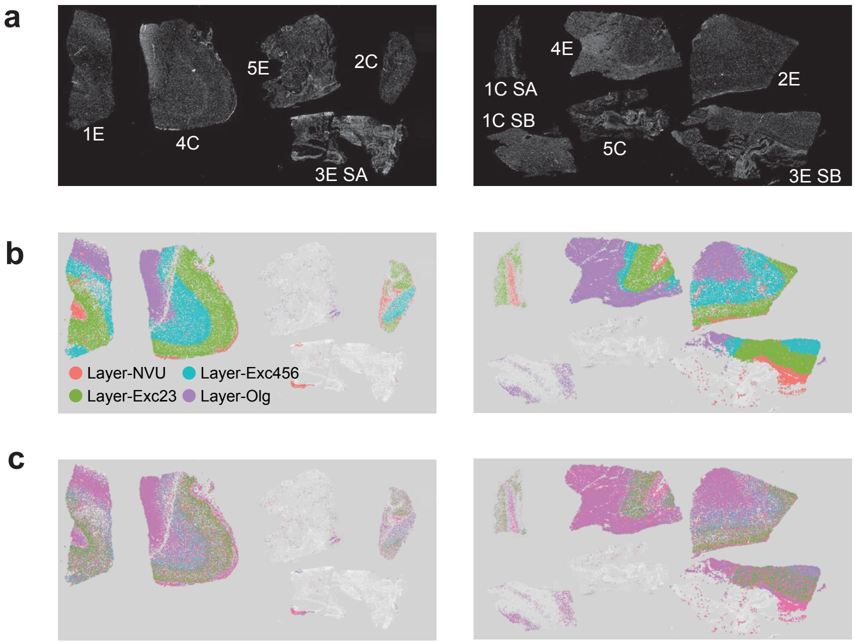
a
1E
4C
5E
2C
3E SA
4E
1C SA
1C SB
5C
2E
3E SB
b
Layer-NVU
Layer-Exc456
Layer-Exc23
Layer-Olg
c

**Extended Data Figure 2. Low resolution images of tissue sections subjected to Xenium spatial transcriptomics**. (a) Images of sections subjected to Xenium spatial analysis, labeled by sample. (b) As in (a) but illustrating identified cellular niches. (c) As in (a) but illustrating different cell types. Legends for colors in (b) and (c) are shown in Figure 1.

**a**

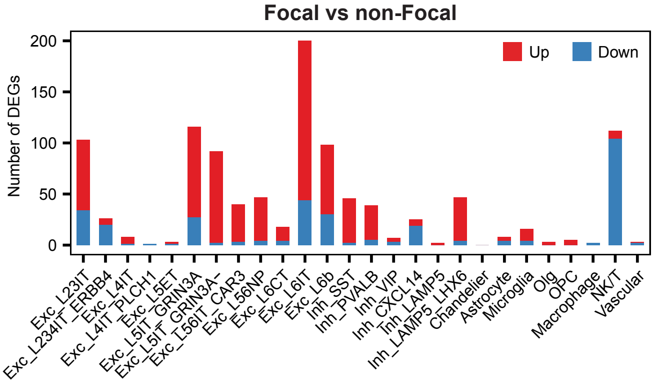

Focal vs non-Focal
Number of DEGs
200
150
100
50
0
Up
Down
Exc_L23IT
Exc_L234IT_ERBB4
Exc_L4IT
Exc_L4IT_PLCH1
Exc_L5ET
Exc_L5IT_GRIN3A
Exc_L5IT_GRIN3A-
Exc_L56IT_CAR3
Exc_L56NP
Exc_L6CT
Exc_L6IT
Exc_L6b
Inh_SST
Inh_PVALB
Inh_VIP
Inh_CXCL14
Inh_LAMP5
Inh_LAMP5_LHX6
Chandelier
Astrocyte
Microglia
Olg
OPC
Macrophage
NK/T
Vascular


**b**

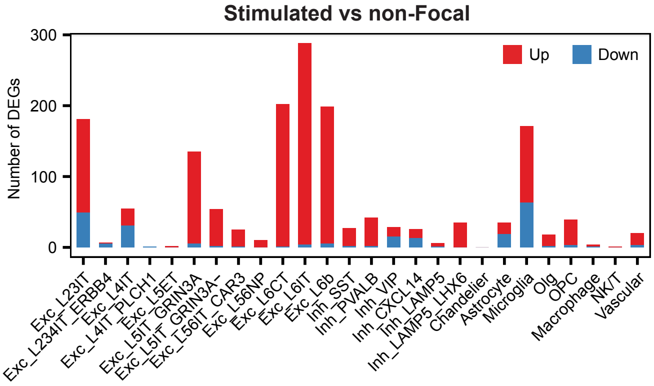

Stimulated vs non-Focal
Number of DEGs
300
200
100
0
Up
Down
Exc_L23IT
Exc_L234IT_ERBB4
Exc_L4IT
Exc_L4IT_PLCH1
Exc_L5ET
Exc_L5IT_GRIN3A
Exc_L5IT_GRIN3A-
Exc_L56IT_CAR3
Exc_L56NP
Exc_L6CT
Exc_L6IT
Exc_L6b
Inh_SST
Inh_PVALB
Inh_VIP
Inh_CXCL14
Inh_LAMP5
Inh_LAMP5_LHX6
Chandelier
Astrocyte
Microglia
Olg
OPC
Macrophage
NK/T
Vascular

**Extended Data Figure 3. Abundance of differentially expressed genes following downsampling**. (a) Numbers of DEGs (y-axis) between focal and non-focal regions across cell types (x-axis). Transcripts upregulated in the focal regions shown in red, downregulated transcripts shown in blue. (b) Same as (a) but illustrating DEGs identified between stimulated and unstimulated non-focal tissue.

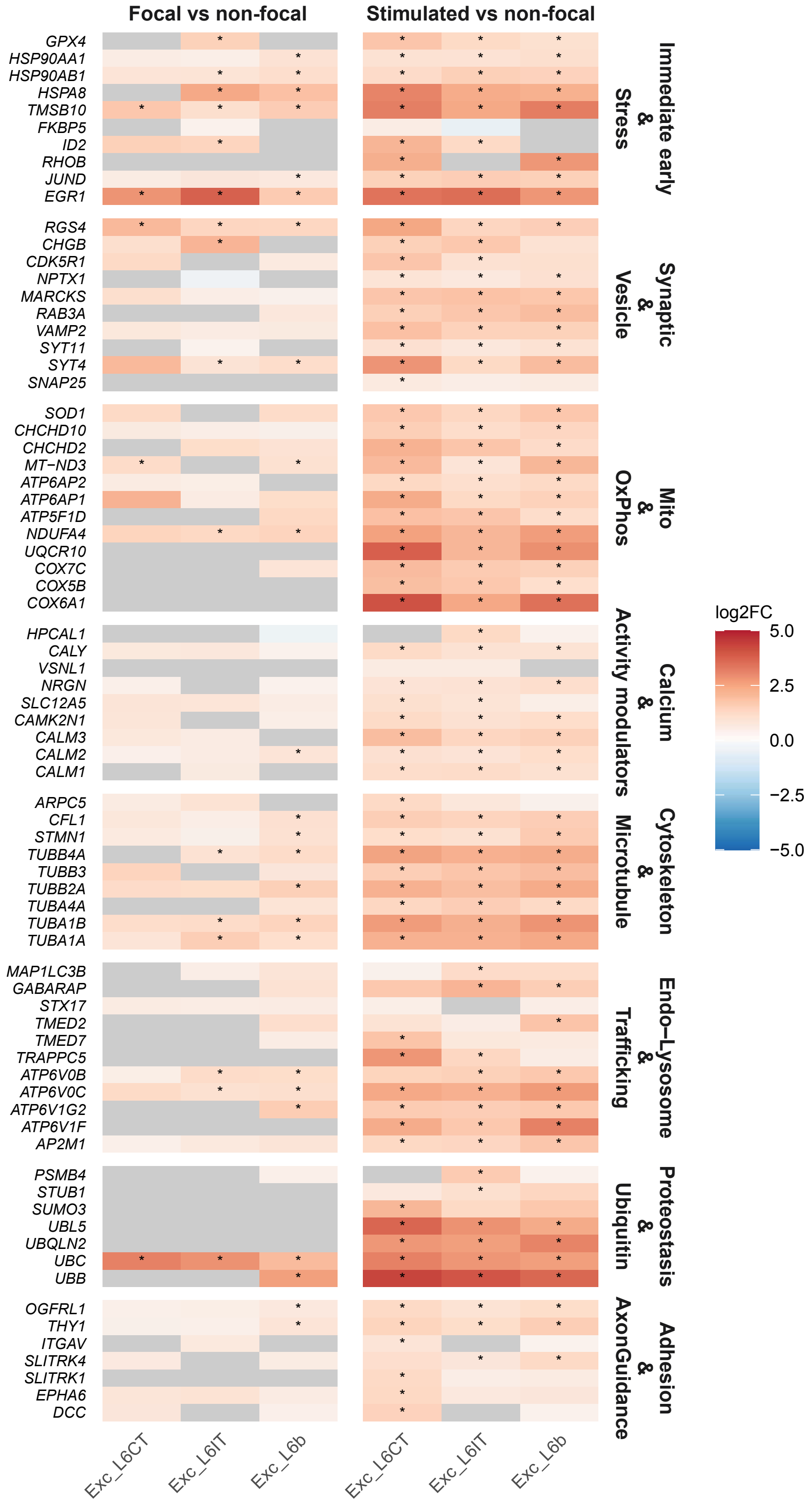

Focal vs non-focal
Stimulated vs non-focal
GPX4
HSP90AA1
HSP90AB1
HSPA8
TMSB10
FKBP5
ID2
RHOB
JUND
EGR1
Immediate early & Stress
RGS4
CHGB
CDK5R1
NPTX1
MARCKS
RAB3A
VAMP2
SYT11
SYT4
SNAP25
Synaptic & Vesicle
SOD1
CHCHD10
CHCHD2
MT−ND3
ATP6AP2
ATP6AP1
ATP5F1D
NDUFA4
UQCR10
COX7C
COX5B
COX6A1
Mito & OxPhos
HPCAL1
CALY
VSNL1
NRGN
SLC12A5
CAMK2N1
CALM3
CALM2
CALM1
Calcium & Activity modulators
ARPC5
CFL1
STMN1
TUBB4A
TUBB3
TUBB2A
TUBA4A
TUBA1B
TUBA1A
Cytoskeleton & Microtubule
MAP1LC3B
GABARAP
STX17
TMED2
TMED7
TRAPPC5
ATP6V0B
ATP6V0C
ATP6V1G2
ATP6V1F
AP2M1
Endo−Lysosome & Trafficking
PSMB4
STUB1
SUMO3
UBL5
UBQLN2
UBC
UBB
Proteostasis & Ubiquitin
OGFRL1
THY1
ITGAV
SLITRK4
SLITRK1
EPHA6
DCC
Adhesion & AxonGuidance
Exc_L6CT
Exc_L6IT
Exc_L6b
Exc_L6CT
Exc_L6IT
Exc_L6b
log2FC
5.0
2.5
0.0
−2.5
−5.0

**Extended Data Figure 4. Comparison of transcripts induced by epileptogenic versus acute activity in layer 6 glutamatergic neurons.** Heatmap showing differential expression of selected gene modules in layer 6 excitatory neuron subtypes (Exc_L6CT, Exc_L6IT, Exc_L6b). The left panel shows focal vs non-focal comparisons, and the right panel shows stimulated vs non-focal comparisons. Genes are grouped into functional categories as listed on the right. Colors represent log2 fold change (log2FC) according to the scale at right, with warmer colors indicating higher expression in the first condition of each comparison. Asterisks denote significant genes ($|\log2FC| > \log2(1.5)$, $FDR < 0.05$) as determined by MAST with patient identity included as a latent variable. Non-significant genes are shown in gray; only genes expressed in at least 10% of cells in either group ($min.pct \geq 0.1$) were tested.

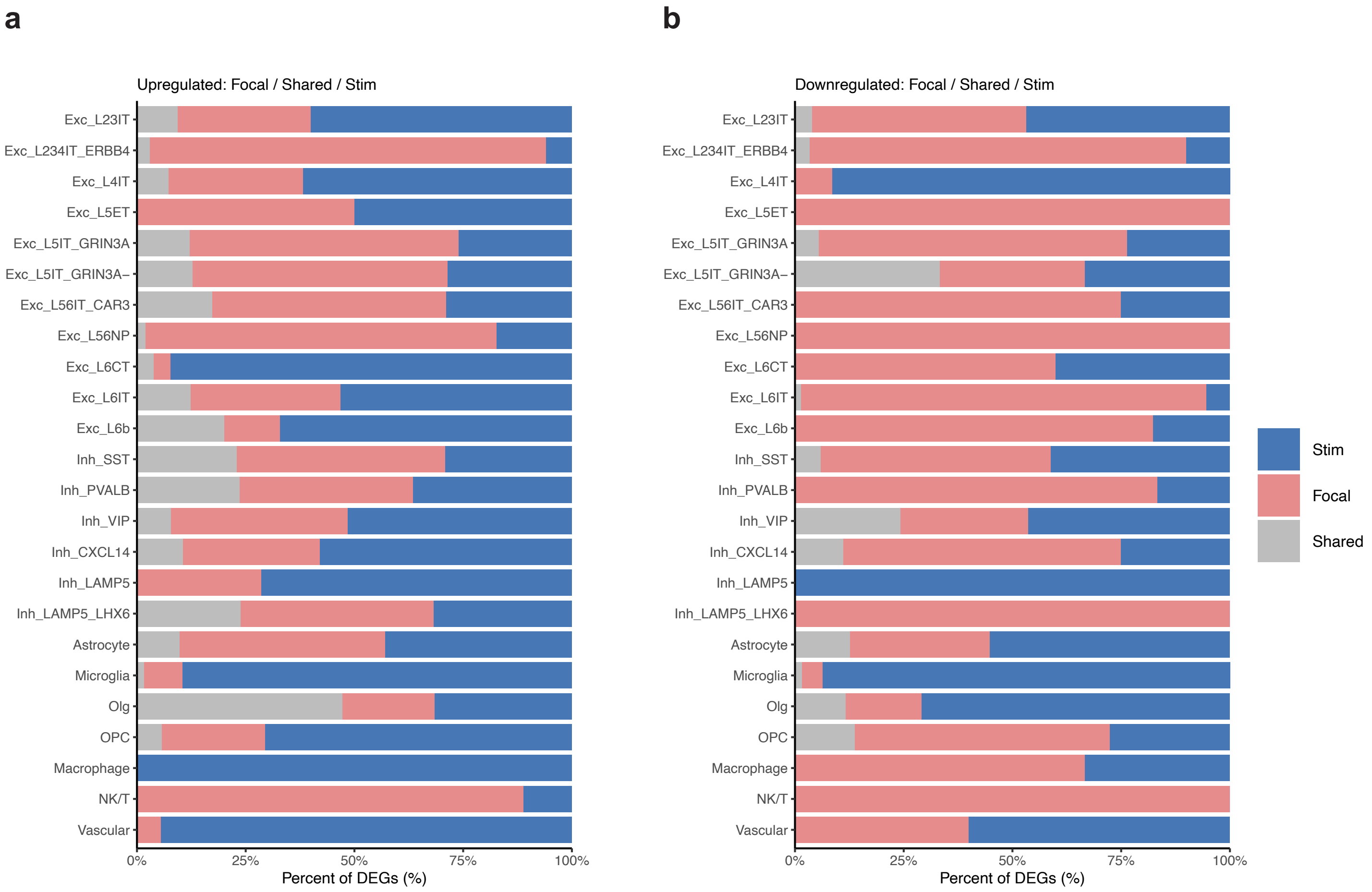
a
Upregulated: Focal / Shared / Stim
b
Downregulated: Focal / Shared / Stim
Exc_L23IT
Exc_L234IT_ERBB4
Exc_L4IT
Exc_L5ET
Exc_L5IT_GRIN3A
Exc_L5IT_GRIN3A−
Exc_L56IT_CAR3
Exc_L56NP
Exc_L6CT
Exc_L6IT
Exc_L6b
Inh_SST
Inh_PVALB
Inh_VIP
Inh_CXCL14
Inh_LAMP5
Inh_LAMP5_LHX6
Astrocyte
Microglia
Olg
OPC
Macrophage
NK/T
Vascular
0%
25%
50%
75%
100%
Percent of DEGs (%)
Stim
Focal
Shared

**Extended Data Figure 5. Overlap between transcripts induced by epileptogenic activity and acute electrode stimulation**. (a) Stacked bar plot illustrating the percentages of DEGs for each cell type that were induced by both epileptogenic and acute activity (gray), only epileptogenic activity (pink), or only acute stimulation (blue). (b) Same as (a) but plotting overlap of downregulated transcripts.

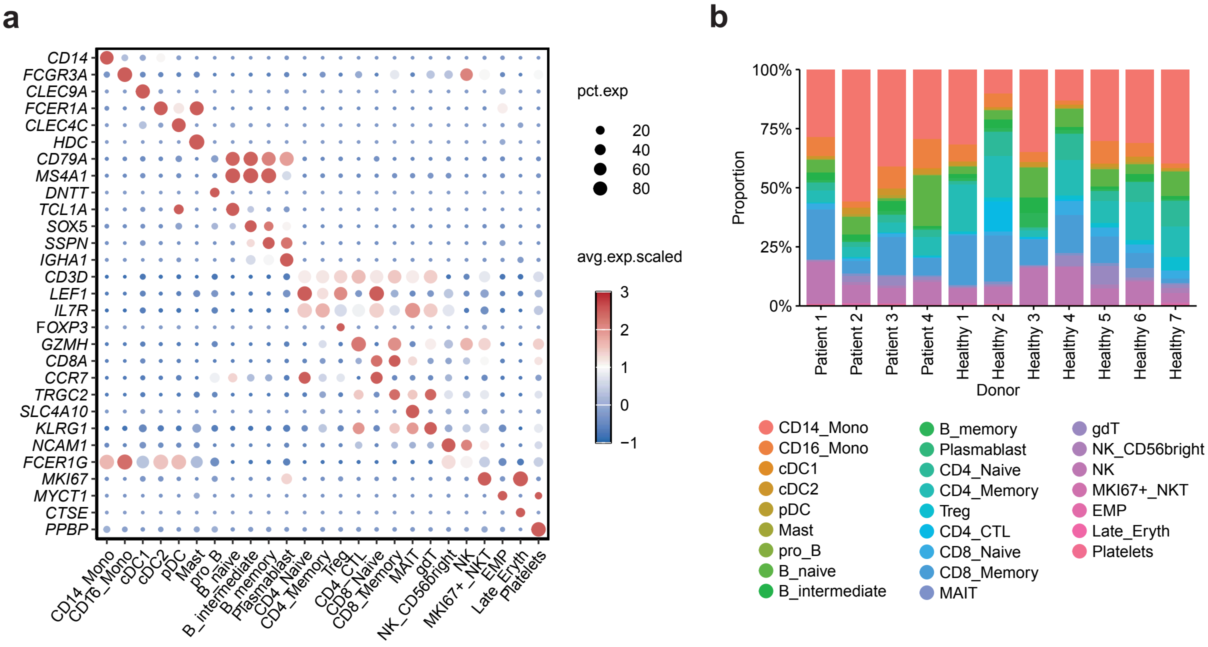
a
CD14
FCGR3A
CLEC9A
FCER1A
CLEC4C
HDC
CD79A
MS4A1
DNTT
TCL1A
SOX5
SSPN
IGHA1
CD3D
LEF1
IL7R
FOXP3
GZMH
CD8A
CCR7
TRGC2
SLC4A10
KLRG1
NCAM1
FCER1G
MKI67
MYCT1
CTSE
PPBP
CD14_Mono
CD16_Mono
cDC1
cDC2
pDC
Mast
pro_B
B_naive
B_intermediate
B_memory
Plasmablast
CD4_Naive
CD4_Memory
Treg
CD4_CTL
CD8_Naive
CD8_Memory
MAIT
gdT
NK_CD56bright
NK
MKI67+_NKT
EMP
Late_Eryth
Platelets
pct.exp
20
40
60
80
avg.exp.scaled
3
2
1
0
−1
b
100%
75%
50%
25%
0%
Proportion
Patient 1
Patient 2
Patient 3
Patient 4
Healthy 1
Healthy 2
Healthy 3
Healthy 4
Healthy 5
Healthy 6
Healthy 7
Donor
CD14_Mono
CD16_Mono
cDC1
cDC2
pDC
Mast
pro_B
B_naive
B_intermediate
B_memory
Plasmablast
CD4_Naive
CD4_Memory
Treg
CD4_CTL
CD8_Naive
CD8_Memory
MAIT
gdT
NK_CD56bright
NK
MKI67+_NKT
EMP
Late_Eryth
Platelets

**Extended Data Figure 6. Annotation and composition of PBMC cell populations across donors.** (a) Dot plot showing expression of canonical marker genes annotated across PBMC populations. Dot size represents the percentage of cells expressing each transcript, and color indicates scaled average expression. (b) Stacked bar plots showing the proportion of each annotated PBMC cell type across individual donors, including patients and healthy controls.
